# Shared genetic effects on chromatin and gene expression reveal widespread enhancer priming in immune response

**DOI:** 10.1101/102392

**Authors:** Kaur Alasoo, Julia Rodrigues, Subhankar Mukhopadhyay, Andrew J. Knights, Alice L. Mann, Kousik Kundu, HIPSCI Consortium, Christine Hale, Gordon Dougan, Daniel J. Gaffney

## Abstract

Noncoding regulatory variants are often highly context-specific, modulating gene expression in a small subset of possible cellular states. Although these genetic effects are likely to play important roles in disease, the molecular mechanisms underlying context-specificity are not well understood. Here, we identify shared quantitative trait loci (QTLs) for chromatin accessibility and gene expression (eQTLs) and show that a large fraction (∼60%) of eQTLs that appear following macrophage immune stimulation alter chromatin accessibility in unstimulated cells, suggesting they perturb enhancer priming. We show that such variants are likely to influence the binding of cell type specific transcription factors (TFs), such as PU.1, which then indirectly alter the binding of stimulus-specific TFs, such as NF-κB or STAT2. Our results imply that, although chromatin accessibility assays are powerful for fine mapping causal noncoding variants, detecting their downstream impact on gene expression will be challenging, requiring profiling of large numbers of stimulated cellular states and timepoints.

---

Genetic differences between individuals can profoundly alter how their immune cells respond to environmental stimuli (1). At the molecular level, these differences manifest as expression quantitative trait loci (eQTLs) that alter the magnitude of gene expression change after stimulation (response eQTLs) (2–7). Although response eQTLs have been implicated in modulating risk for complex immune-mediated disorders (8,9), the molecular mechanisms that give rise to these context specific effects are poorly understood. The majority of eQTLs also alter chromatin accessibility, presumably reflecting disruption of transcription factor (TF) binding (10). Because cellular response to external stimuli is regulated by stimulus-specific transcription factors (TFs), response eQTLs might directly disrupt their binding (Fig. 1A). In support of this model, a number of studies have observed that response eQTLs are enriched at the binding sites of stimulation-specific TFs such as NF-κB and STAT2 (5–7). However, a single stimulus or a developmental cue can upregulate alternate sets of genes in different cell types, even when the activated signalling pathways and TFs remain the same (11). To explain these observations, multiple studies have proposed a hierarchical enhancer activation model (11–14), under which cell type specific TFs bind to a subset of enhancers without a direct effect on target gene expression. This enhancer ‘priming’ can facilitate their subsequent activation by signal specific TFs, producing a cell type specific response (Fig. 1B). Thus, genetic variants could modulate stimulus specific effects on gene expression *indirectly*, by altering the binding of a cell type specific TF, for example PU.1 in macrophages, that regulate chromatin accessibility (Fig 1B). However, the genome-wide prevalence of enhancer priming is currently unclear because directed genome editing studies have been limited to handful of loci (15, 16). A powerful alternative is to use shared genetic associations at chromatin and gene expression level to probe the relationships between enhancer accessibility and gene transcription.

**Fig. 1.**
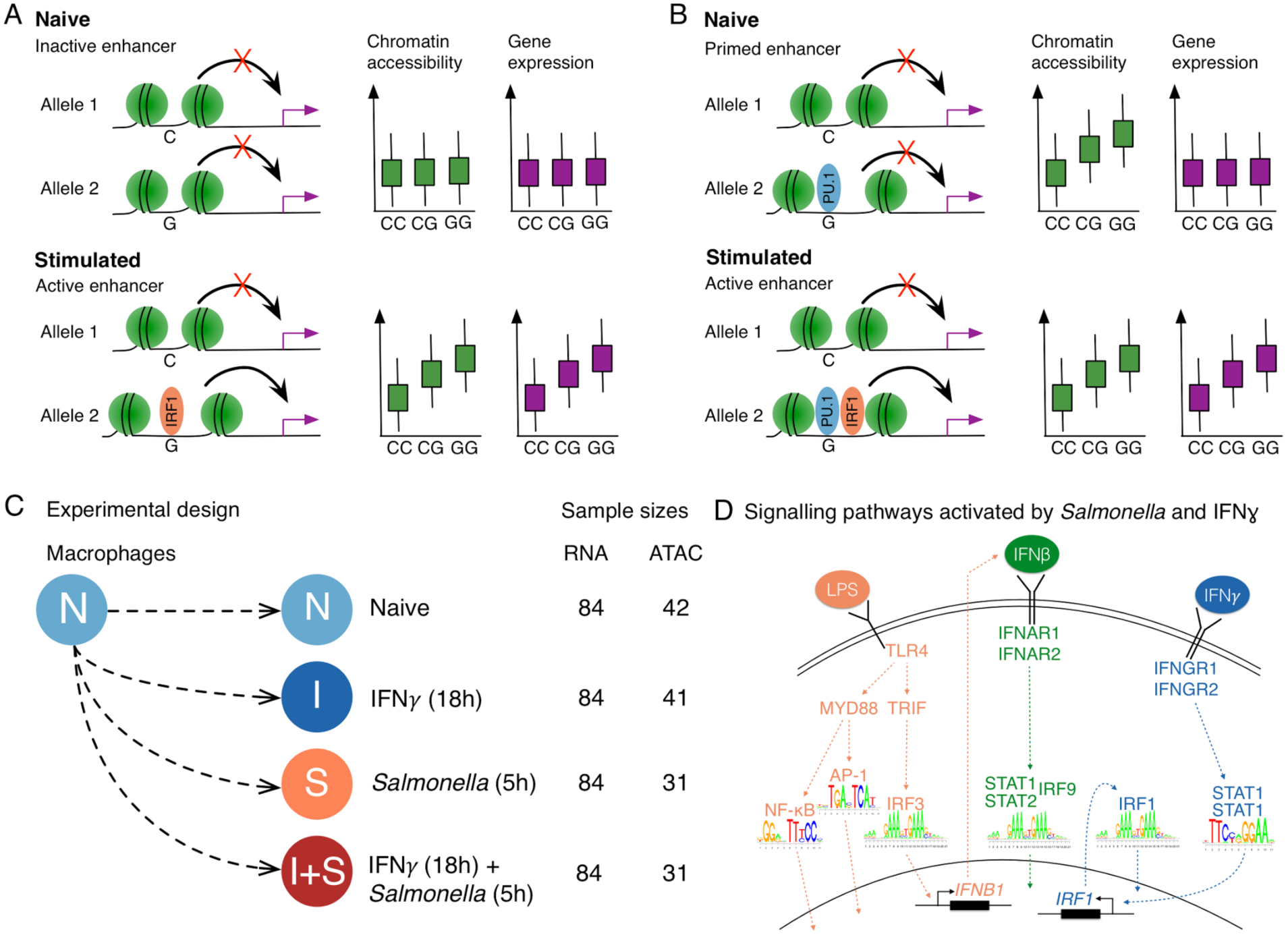
Regulation of gene expression in macrophage immune response. (**A**) Genetic variant has a direct effect on the binding of a stimulation-specific TF (IRF1) and target gene activation. (**B**) Genetic variant in a primed enhancer disrupts the binding of a cell type specific TF (e.g. PU.1) that indirectly influences stimulation-specific TF (IRF1) binding via modulation of chromatin accessibility. (**C**) Overview of the experimental design. (**D**) TLR4 recognises lipopolysaccharide (LPS) on *Salmonella* cell wall and activates NF-κB, AP-1 and IRF3 transcription factors (TFs) (21). IRF3 stimulates IFNβ production that culminates with the activation of STAT1-STAT2-IRF9 complex. IFNɣ binds to IFNɣ receptor and activates STAT1 and IRF1 TFs (22).

We focussed on enhancer priming in the context of human macrophage immune response. To ensure sufficient numbers of cells, we differentiated macrophages from a panel of 123 human induced pluripotent cell lines (iPSCs) obtained from the HipSci project (17, 18). We profiled gene expression (RNA-seq) and chromatin accessibility (ATAC-seq) in a subset of 86 successfully differentiated lines (fig. S1, table S1) in four experimental conditions: naive (N), 18 hours IFNɣ stimulation (I), 5 hours *Salmonella enterica* serovar Typhimurium (*Salmonella*) infection (S), and IFNɣ stimulation followed by *Salmonella* infection (I+S) (Fig. 1C). We chose these stimuli because they activate distinct, well characterised signalling pathways (Fig. 1D, fig. S2) and pre-stimulating macrophages with IFNɣ prior to bacterial infection is known to lead to enhanced microbial killing and stronger activation of the inflammatory response (19, 20).

We identified common genetic variants that were associated with either gene expression (eQTLs) or chromatin accessibility (caQTLs). Using an allele-specific method implemented in RASQUAL (23), we detected at least one QTL for up to 3,431 genes and 20,788 chromatin regions (caQTL regions) in each condition (10% FDR) (fig. S3), 50-75% of which were shared between conditions (fig. S3). Next, using a statistical interaction test followed by filtering on effect size, we identified 387 response eQTLs and 2247 response caQTLs with a small or undetectable effect (fold change < 1.5) in the naive state that increased at least 1.5 fold after stimulation (see Methods). These genetic effects displayed a variety of activity patterns (Fig. 2A, fig. S4). Strikingly, 18% of the response eQTLs appeared only after the cells were exposed to both stimuli (cluster 1), exceeding the number that appeared after IFNɣ stimulation alone (clusters 5 and 6). Response caQTL regions harboured closed chromatin in the naive cells (median transcripts per million (TPM) = 0.49) and became 3.8-fold more accessible only after the relevant stimulus (fig. S4). Furthermore, response caQTLs were enriched for disrupting stimulus-specific TF motifs (fig. S4), suggesting that they are largely driven by TFs that bind to DNA only after stimulation.

**Fig. 2.**
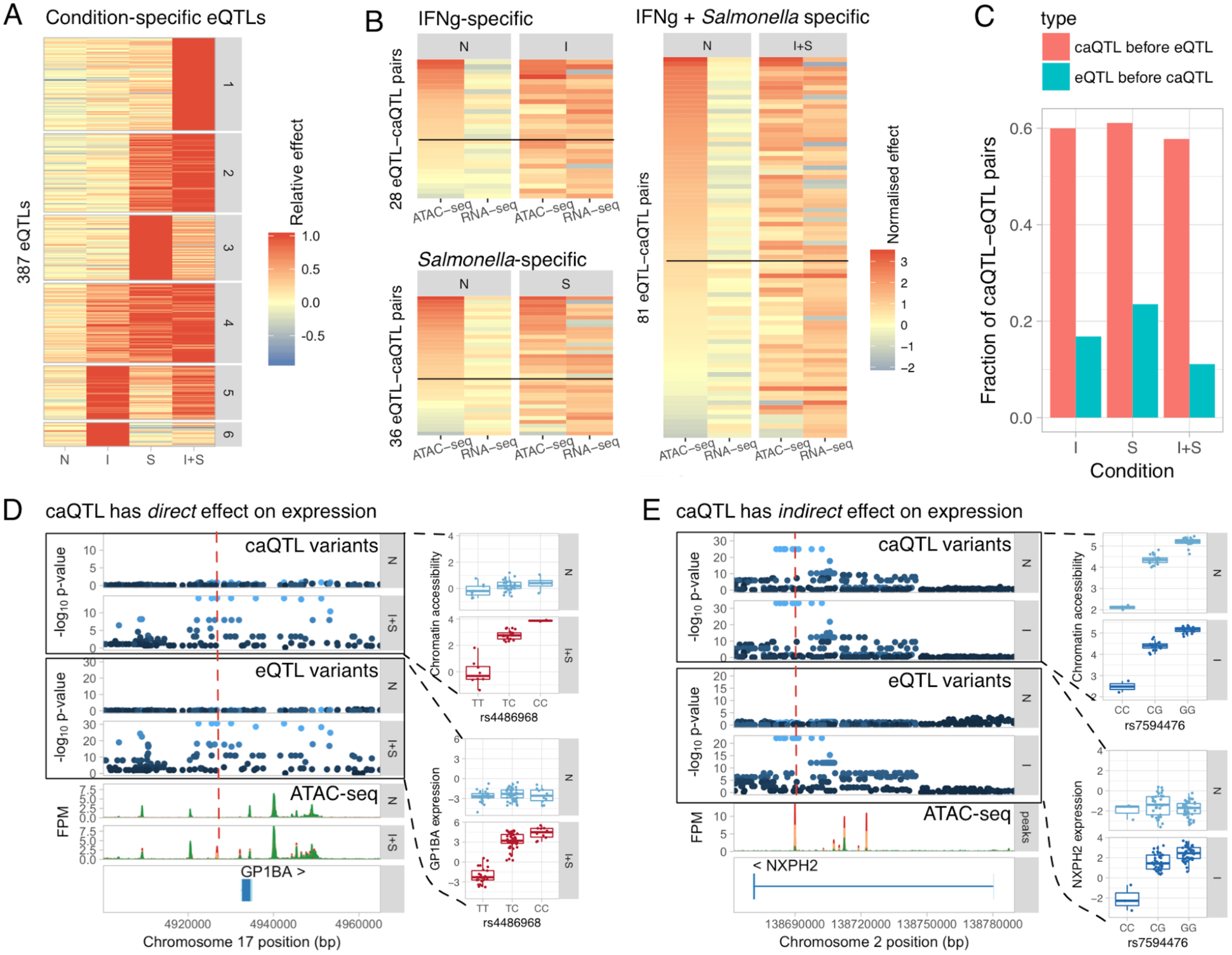
Quantifying the extent of enhancer priming in macrophage immune response. **(A)**Clustering the effect sizes of response eQTLs. Response eQTLs appear after IFNɣ stimulation (clusters 5 and 6), *Salmonella* infection (clusters 2-4) or only when both of the stimuli are present (cluster 1). **(B)** Effect sizes of eQTL-caQTL pairs in naive and stimulated conditions. The pairs are grouped by the condition in which the eQTL had the largest effect size (I, S or I+S). The heatmaps are sorted by caQTL effect size in the naive condition (first column). Approximately 60% of the caQTLs linked to response eQTLs are present in the naive condition (fold change > 1.5, above solid black line). **(C)** Comparison of our estimated rate of enhancer priming (caQTL precedes response eQTL) to a negative control (eQTL precedes a response caQTL). **(D)** The rs4486968 variant (red dashed line) becomes simultaneously associated with chromatin accessibility and GP1BA expression after IFNɣ + *Salmonella* stimulation. **(E)** A genetic variant (red dashed line) regulates chromatin accessibility in the naive cells and becomes associated with NXPH2 expression only after IFNɣ stimulation. FPM, fragments per million.

To quantify the extent of enhancer priming in macrophage immune response, we next focussed on how response eQTLs manifest on the chromatin level. We grouped response eQTLs (Fig. 2A) by the condition in which they had the largest effect size (I, S or I+S). We then used linkage disequilibrium (LD) (R^2^ > 0.8) between the lead variants to identify caQTL-eQTL pairs that were likely to be driven by the same causal variant (see Methods). For example, we identified a QTL upstream of GP1BA that had no effect in naive cells, but became simultaneously associated with chromatin accessibility and gene expression after IFNɣ + *Salmonella* stimulation (Fig. 2D). The lead caQTL variant (rs4486968) was predicted to disrupt NF-κB binding motif (fig. S5), illustrating how a genetic variant can have direct effect on stimulus-specific TF binding and gene expression. In contrast, a genetic variant in an intron of NXPH2 modulated the accessibility of a regulatory element both in naive and stimulated cells, but only became associated with gene expression after IFNɣ stimulation (Fig. 2E). Genome-wide, we found that for approximately half of all response eQTLs, the linked caQTL was present in naive cells prior to stimulation (caQTL fold change > 1.5), suggesting that many response eQTLs disrupt enhancer priming (Fig. 2B).

One potential issue with our analysis is that using LD to identify eQTL-caQTL pairs will sometimes lead to false positives where two independent causal variants for different phenotypes are mistaken for a single shared causal variant. To estimate our false positive rate, we performed a reverse analysis where we asked how often response caQTLs were linked to eQTLs that were present in the naive state, reasoning that these are likely to be false positives. Using the same fold change threshold as above, we estimated the mean false positive rate to be 17% (Fig. 2C).

We speculated that response eQTLs that alter enhancer priming should be enriched for disrupting the motifs of macrophage cell type specific TFs. To test this, we focussed on the 145 eQTL-caQTL pairs (137 unique caQTLs) identified above (Fig. 2B). We found that 9/78 caQTLs present in the naive cells disrupted PU.1 motifs compared to none of the 59 caQTLs that appeared together with the response eQTL (Fisher’s exact test, p = 0.01). For example, the rs7594476 variant in the NXPH2 enhancer disrupted PU.1 binding in a direction consistent with the caQTL effect (Fig. 3A).

**Fig. 3.**
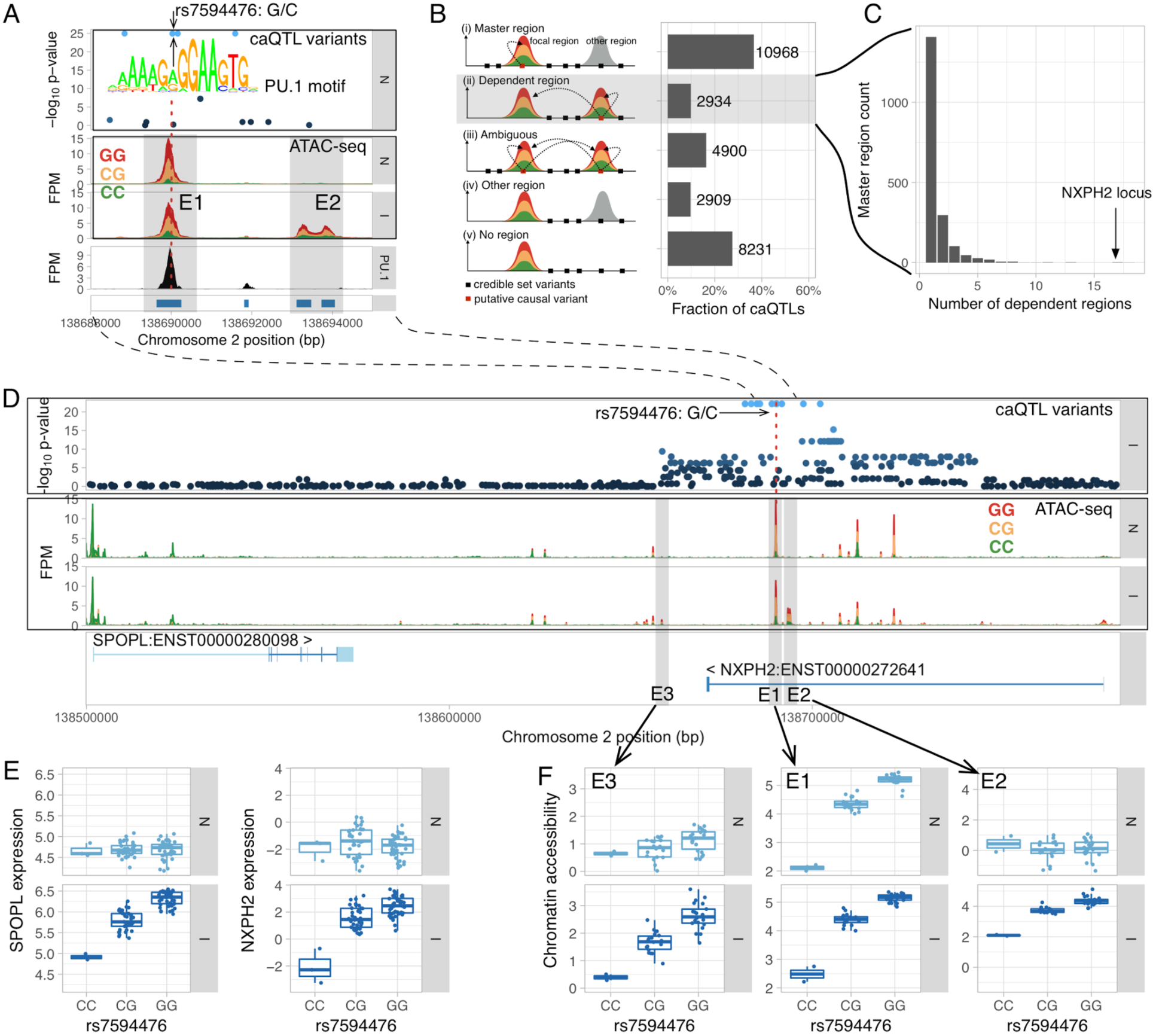
Identifying caQTLs that regulate chromatin accessibility at multiple independent regions. (**A**) Fine mapping the putative causal variant at the NXPH2 locus. The master caQTL region (E1) overlaps a PU.1 ChIP-seq peak from (27). Only one of the two variants (rs7594476) within the E1 region is predicted to disrupt the PU.1 motif. No associated variants overlap the IFNɣ-specific dependent region E2. (**B**) Classifying caQTLs into master regions (i), dependent regions (ii) and ambiguous cases where the credible set overlaps either multiple regulated regions (iii) or does not overlap any regulated regions (iv-v). (**C**) Histogram of the number of associated dependent regions for each master region. (**D**) Multiple open chromatin regions regulated by a single caQTL at the NXPH2 locus. Master caQTL region (E1) and two IFNɣ-specific dependent regions (E2 and E3) are highlighted by grey shadows. FPM, fragments per million. (**E**) Master caQTL variant (red dashed line on panel D) becomes associated with NXPH2 and SPOPL expression after IFNɣ stimulation. (**F**) Boxplots of the master caQTL region (E1) and two dependent regions (E2 and E3) stratified by the genotype of the caQTL lead variant.

Recent evidence suggests that single genetic variants can modulate the activity of multiple regulatory elements within topologically associated domains (23–26). One plausible mechanism for these broad associations is that a single causal variant may directly regulate the accessibility of a “master” region, which subsequently influences neighbouring “dependent” regions (23). We used caQTL summary statistics to heuristically identify likely master and dependent regions, assuming that the causal variant should reside within the master region itself, and this affects accessibility in dependent regions (Fig. 3B) (see Methods). We found a striking example of such a relationship at the NXPH2 locus, where a putative causal variant in the master region was also associated with the accessibility of neighbouring dependent region after IFNɣ stimulation (Fig. 3A). Using this approach, we identified 2,934 dependent regions that belonged to 1,921 unique master regions (Fig. 3B). While 77% of the master regions had a single dependent region only a few kb away (fig. S6), we found many loci where master peaks were associated with multiple regions of open chromatin (Fig. 3C). In the NXPH2 locus introduced above, we detected 18 dependent regions spanning 100 kilobases of DNA (Fig. 3C), six of which appeared only after IFNɣ stimulation (Fig. 3D,F). Notably, the appearance of condition-specific dependent regions correlated with the caQTL becoming a response eQTL for both NXPH2 and SPOPL (Fig. 3E), suggesting that some of them might be required for gene activation. Using a linear model followed by strict filtering (see Methods), we found a total of 64 condition-specific dependent regions genome-wide, two of which are highlighted in fig. S7.

Because they can be engineered with high efficiency, iPSC-derived cells are promising cellular models of disease. Similarly to previous studies (7), we found that macrophage eQTLs and caQTLs were enriched for GWAS hits of multiple immune-mediated disorders (fig. S8).

However, observing a genome-wide enrichment has only limited utility and detailed follow up of a locus is only justified when there is evidence for a shared causal mechanism between GWAS and eQTL associations. Thus, we used a statistical colocalisation test *(28)* to identify cases where the gene expression and trait association signals were consistent with a model of a single, shared causal variant. We identified 22 eQTLs (table S3) that showed evidence of colocalisation (PP3 + PP4 > 0.8, PP4/PP3 > 9) with at least one disease (see Methods).

Consistent with our enrichment analysis, we found the largest number of overlaps with IBD and RA (Fig. 4A). Interestingly, only 10/22 of the colocalised eQTLs were detected in the naive cells and each additional stimulated state increased the number of overlaps by approximately 30% (Fig. 4B). For example, we found an IFNɣ + *Salmonella* specific response eQTL for TRAF1 that colocalised with a RA GWAS hit (fig. S9). Although the same overlap was previously reported in whole blood (29), our data highlights the environmental condition in which the association is active.

**Fig. 4.**
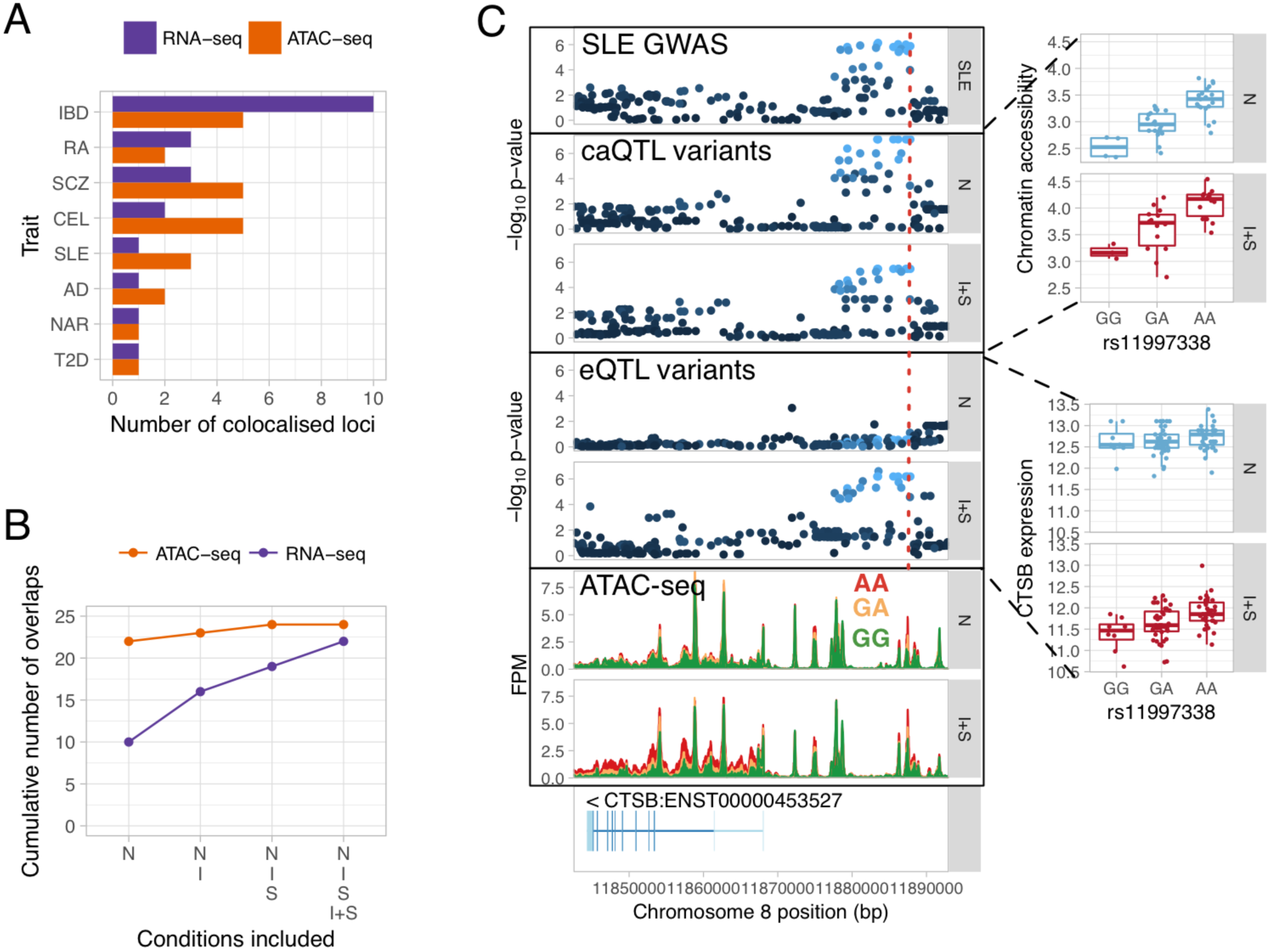
Identifying eQTLs and caQTLs that colocalise with complex disease risk loci. **(A)** Total number of colocalised GWAS hits identified for each trait across the four conditions. **(B)** Cumulative number of colocalised GWAS hits identified by starting with overlaps in the naive condition and sequentially adding IFNɣ, *Salmonella* and IFNɣ + *Salmonella* conditions. **(C)** Constitutive caQTL and IFNɣ + *Salmonella* specific eQTL for CTSB colocalised with a GWAS association for systemic lupus erythematosus (SLE). FPM, fragments per million. Disease acronyms: IBD, inflammatory bowel disease; RA, rheumatoid arthritis; SLE, systemic lupus erythematosus; AD, Alzheimer's disease; SCZ, schizophrenia; T2D, type 2 diabetes; NAR, narcolepsy; CEL, celiac disease.

Our analysis of enhancer priming suggested that many disease associations might manifest at the level of chromatin without an apparent effect on expression. To explore this further, we focussed on colocalisation between caQTLs and GWAS hits. We detected 24 caQTLs that colocalised with a GWAS hit (table S4), but only two of these also colocalised with an eQTL (PTK2B eQTL with Alzheimer's disease (fig. S10) and WFS1 eQTL with type 2 diabetes). Since genes often have multiple independent eQTLs (30), we reasoned that some caQTLs might be secondary eQTLs for their target genes. To capture these secondary effects, we first identified four additional genes that were associated with a caQTL lead variant at FDR < 10%, even though the caQTL and eQTL lead variants were not in strong LD (i.e. R^2^ < 0.8). We repeated the colocalisation analysis on these loci and identified two additional overlaps (table S3), including a secondary eQTL for CTSB that colocalised with a GWAS hit for systemic lupus erythematosus (SLE) (Fig. 4C). Interestingly, although the CTSB eQTL appeared after IFNɣ + *Salmonella* stimulation, the caQTL was already present in naive cells. Although some caQTL colocalisation with eQTLs might remain undetected due to lack of power, the CTSB example suggests that a fraction of disease-associated caQTLs might correspond to primed enhancers that regulate gene expression in some other yet unknown conditions. Although majority (22/24) of caQTL overlaps with disease were detected in the naive cells (Fig. 4C), this is confounded with a smaller ATAC-seq sample size in *Salmonella* and IFNɣ + *Salmonella* conditions that limited our power to detect colocalisations.

## Discussion

The results of our study resolve an apparent paradox in the genetics of human complex traits: although disease loci from association studies are strongly enriched in regulatory elements (31, 32), a relatively small fraction are explained by known eQTLs, even those identified in trait-relevant tissues (29, 33, 34). Our results suggest that this apparent contradiction arises partly because many disease risk variants affect chromatin structure in a broad range of cellular states, but their impact on expression is highly context-specific. This conclusion is supported by studies of 3D chromatin structure linking GWAS loci to putative target genes but with no observable effect on gene expression (35), in particular because enhancer-promoter interactions are known to precede transcription (36, 37). We believe our result has important implications for future studies of human disease. First, it is likely that a large range of cellular states will need to be profiled in order to capture the effects of disease-associated variants on expression. Our results suggest this space will be challenging to explore systematically, especially given the numbers of novel associations we detect using a combination of just two stimuli. Second, overlap of disease variants with open chromatin, while likely to be informative regarding the identity of the causal variant, may be less useful predictors of the disease relevant cell state.

Although our study suggests that many human disease associated variants impact enhancer priming, the functional relevance of this is currently not well understood. First, enhancer priming may facilitate cell type specific response to ubiquitous signals (11, 38, 39). Although specificity can also be achieved by cooperative binding to newly established enhancers (40), TFs differ in their intrinsic ability to bind to closed chromatin (41). Thus, enhancer priming might be a preferred mechanism of cooperation between ‘pioneer’ TFs that can independently open up chromatin (e.g. PU.1 in macrophages) and ‘settlers’ (e.g. NF-κB) that predominantly bind to accessible regions (42). Alternatively, enhancer priming might facilitate rapid response to external stimuli. In support of this model, promoters of immediate early response genes are already accessible in naive cells (43) and TF binding to primed enhancers peaks minutes after stimulation while the activation of *de novo* enhancers can take several hours (40). Thus, response eQTLs that appear rapidly after stimulation might be enriched for primed enhancers relative to those that appear later. Finally, enhancer priming might not be limited to single regulatory elements. Our results (Fig. 3D) together with previous reports (16, 44) suggest that some regulatory elements can act as ‘seed’ enhancers that allow other neighbouring enhancers to become active after stimulation and lead to upregulation of gene expression. Although we have identified a small number of such examples, future caQTL mapping studies in multiple cell types and conditions have a potential to systematically identify and characterise these hierarchical relationships between enhancers.

In summary, our results illustrate how pre-existing genetic effects on chromatin propagate to gene expression during immune activation, and highlights the relevance of these hidden genetic effects for deciphering the molecular architecture of disease-associated variants. Our study is also the first that we are aware of to utilise iPSC-derived cells to study genetic effects in immune response. We believe a major future use of this system will be the systematic exploration of gene-environment interactions across large numbers of cell states. Furthermore, because iPSCs are readily engineered, the identity of causal variants and their downstream consequences can be directly tested in exactly the same cell types and conditions where they were discovered.

## Acknowledgements

We thank Leopold Parts, Jeremy Schwartzentruber, Chris Wallace, Lili Milani, Kaido Lepik and Hedi Peterson for helpful comments on the manuscript. We thank Rachel Nelson for assistance and early access to HipSci iPSC lines. We also thank WTSI DNA Pipelines and Cytometry Core Facility for their sequencing and flow cytometry services. This work was supported by the Wellcome Trust grant #098051. K.A. was supported by a PhD fellowship from the Mathematical Genomics and Medicine programme from the Wellcome Trust. The iPSC lines were generated at the Wellcome Trust Sanger Institute, under the Human Induced Pluripotent Stem Cell Initiative funded by a strategic award (WT098503) from the Wellcome Trust and Medical Research Council. We also acknowledge Life Science Technologies Corporation as the provider of cytotune.

## Supplementary Materials

### Cell culture and reagents

#### Donors and cell lines

Human induced pluripotent stem cells (iPSCs) from 123 healthy donors (72 female and 51 male) (table S1) were obtained from the HipSci project (18). Of these lines, 57 were initially grown in feeder-dependent medium and 66 were grown in feeder-free E8 medium.

#### Feeder-free iPSC culture

Feeder-free iPSCs were grown on tissue culture treated plates coated with vitronectin (VTN-N) (Gibco, cat. no. A14700) in Essential 8 (E8) medium (Gibco). The cells were dissociated from the plates using Gentle Cell Dissociation Buffer (Stemcell Technologies, cat. no. 07174) and passaged every 3-5 days. Prior to macrophage differentiation, the feeder-free iPSCs were first transferred to feeder-dependent media and propagated for at least two passages. This step was necessary because multiple attempts to differentiate macrophage directly from feeder-free iPSCs with our protocol failed.

#### Feeder-dependent iPSC culture

Feeder-dependent iPSCs were grown on irradiated CF-1 mouse embryonic fibroblast (MEF) feeder cells (AMS Biotechnology) in Advanced DMEM-F12 (Gibco) supplemented with 20% Knock-Out Serum Replacement (KSR) (Gibco), 2mM L-glutamine (Sigma), 50 IU/ml penicillin (Sigma), 50 IU/ml Streptomycin (Sigma) and 50µM β-Mercaptoethanol (Sigma M6250). The media was supplemented with 4 ng/ml recombinant human fibroblast growth factor (rhFGF) basic (R&D, 233-FB-025) to maintain pluripotency and was changed daily. MEFs were seeded on 0.1% gelatine-coated tissue-culture treated plates (Corning 6-well or 10 cm plates) 24 hours prior to passaging iPSCs at a cell density of 2 million cells/6-well or 10-cm plate in Advanced DMEM-F12 supplemented with 10% FBS (labtech), 2mM L-glutamine (Sigma), 50 IU/ml Penicillin and 50 IU/ml Streptomycin (Sigma). Prior to passaging or embryoid body formation, iPSCs were dissociated from the plates using 1:1 mixture of collagenase (1 mg/ml) and dispase (1 mg/ml) (both Gibco).

#### Macrophage differentiation protocol

iPSCs were differentiated into macrophages using a previously published protocol (45) involving 3 stages: i) embryoid body (EB) formation, ii) generation of monocyte-like myeloid progenitors from the EBs and iii) terminal differentiation of the progenitors into macrophages. For EB formation, iPSC colonies were treated with 1:1 mixture of collagenase (1 mg/ml) and dispase (1 mg/ml) and intact colonies were transferred to low-adherence plates (Sterilin). The colonies were cultured in feeder-dependent iPSC medium without rhFGF for 3 days. On day 3, the EBs were harvested and transferred to gelatinised tissue-culture treated 10 cm plates in serum-free haematopoietic medium (Lonza X-VIVO 15), supplemented with 2mM L-glutamine (Sigma), 50 IU/ml penicillin, 50 IU/ml streptomycin (Sigma), 50µM β-Mercaptoethanol (Sigma M6250), 50 ng/ml macrophage colony stimulating factor (M-CSF) (R&D) and 25 ng/ml interleukin-3 (IL-3) (R&D). EBs were maintained in these plates with media changes every 3-5 days for 4-6 weeks until the progenitor cells appeared in the supernatant. Progenitor cells were harvested from the supernatant, filtered through a 40µM cell strainer (BD 352340), centrifuged at 1200 rpm for 5 minutes, counted, and plated in RPMI 1640 (Gibco) supplemented with 10% FBS (labtech), 2mM L-glutamine (Sigma) and 100 ng/ml hM-CSF (R&D) at a cell density of 150,000 cells per well on a 6-well plate and differentiated for another 7 days.

#### Differentiation outcomes

We performed 138 macrophage differentiation attempts from 123 distinct HipSci iPSC lines (table S1). We were able to differentiate macrophages from 101/123 (82%) of the iPSC lines. Successful differentiation means that we obtained at least some proportion of cells that exhibited characteristic spindle-like macrophage morphology. For 97/101 lines, we further confirmed the cell surface expression of CD14, CD16 and CD206 macrophage markers using flow cytometry. However, some of the differentiated lines did not produce enough macrophages to perform all of the experimental assays or the differentiated cells were not pure enough to be used in stimulation experiments. In total, we obtained high quality RNA-seq data from 89 differentiations corresponding to 85 unique donors and ATAC-seq data from up to 42 unique donors in up to four experimental conditions (table S1).

### *Salmonella* infection and IFNɣ stimulation

Two wells of a 6-well plate were used per condition to ensure sufficient amount of RNA. On day 6 of macrophage differentiation, medium was changed for all wells with half of the wells receiving macrophage differentiation media (with M-CSF) and the other half of the cells receiving macrophage differentiation media supplemented with 20 ng/ml IFNɣ (R&D) and M-CSF. After 18 hours, cells from two wells of the naive and IFNɣ conditions were harvested for RNA extraction. The remaining two wells from each condition were additionally infected with *Salmonella enterica* serovar Typhimurium SL1344 (hereafter *Salmonella*) for 5 hours.

Two days before infection, *Salmonella* culture was inoculated in 10 ml low salt LB broth and incubated overnight in a shaking incubator (200 rpm) at 37°C. Next morning, the culture was diluted 1:100 into 10 ml of fresh LB broth and incubated again in a shaking incubator. In the afternoon the culture was diluted once more 1:100 into 45 ml of LB broth and kept overnight in a static incubator. In the morning before infection, the culture was centrifuged at 4000 rpm for 10 minutes, washed once with 4°C PBS and resuspended in 30 ml of PBS. Subsequently, optical density at 600 nm was measured and *Salmonella* was diluted in macrophage differentiation media (without MCSF) at multiplicity of infection (MOI) 10 assuming 300,000 cells per well. To infect the cells, old media was removed and replaced with 1 ml of media containing *Salmonella* for 45 minutes. Subsequently, the cells were washed twice with PBS and replaced in fresh medium with 50 ng/ml gentamicin (Sigma) to kill extracellular bacteria. After 45 minutes the medium was changed once again to fresh medium containing 10 ng/ml gentamicin.

For RNA extraction, cells were washed once with PBS and lysed in 300 ul of RLT buffer (Qiagen) per one well of a 6-well plate. Lysates from two wells were immediately pooled and stored at -80°C. RNA was extracted using RNA Mini Kit (Qiagen) following manufacturer's instructions and eluted in 35 µl nuclease-free water. RNA concentration was measured using NanoDrop and RNA integrity was measured on Agilent 2100 Bioanalyzer using RNA 6000 Nano total RNA kit.

### Flow cytometry

We used flow cytometry to measure the cell surface expression of three canonical macrophage markers: CD14, CD16 (FCGR3A/FCGR3B) and CD206 (MRC1). Macrophages were cultured in 10 cm tissue-culture treated plates and detached from the plates by incubation in 6 mg/ml lidocaine-PBS solution (Sigma L5647) for 30 minutes followed by gentle scraping. From each cell line we harvested between 300,000-500,000 cells. Detached cells were washed in media, centrifuged at 1200 rpm for 5 minutes and resuspended in flow cytometry buffer (2% BSA, 0.001% EDTA in D-PBS) and split into two wells of a 96-well plate. Nonspecific antibody binding sites were blocked by incubating cells with Human TruStain FcX (Biolegend) for 45 minutes and washing with flow cytometry buffer. Half of the cells were stained for 1 hour with the PE-isotype control (BD 555749) antibody. The other half of the cells were co-stained for 1 hour with following three antibodies: CD14-Pacific Blue (BD 558121), CD16-PE (BD 555407), CD206-APC (BD 550889). After staining, the cells were washed three times. Resuspended cells were filtered through cell-strainer cap tubes (BD 352235) and measured on the BD LSRFortessa Cell Analyzer. The raw flow cytometry data has been deposited to Zenodo (doi: 10.5281/zenodo.234214).

### RNA sequencing

All of the RNA-seq libraries were constructed using poly-A selection. The first 120 RNA-seq libraries from 30 donors were constructed manually using the Illumina TruSeq stranded library preparation kit. The TruSeq libraries were quantified using Bioanalyzer and manually pooled for sequencing. For the remaining 216 samples, we used an automated library construction protocol that was based on the KAPA stranded mRNA-seq kit. The KAPA libraries were quantified using Quant-iT plate reader and pooled automatically using the Beckman Coulter NX-8. The first 16 samples were sequenced on Illumina HiSeq 2500 using V3 chemistry and multiplexed at 4 samples/lane. All of the other samples were sequenced on Illumina HiSeq 2000 using V4 chemistry and multiplexed at 6 samples/lane. Sample metadata is presented in table S5.

#### RNA-seq preprocessing and quality control

RNA-seq reads were aligned to the GRCh38 reference genome and Ensembl 79 transcript annotations using STAR v2.4.0j (46). Subsequently, VerifyBamID v1.1.2 (47) was used to detect and correct any potential sample swaps and cross-contamination between donors. We did not detect any cross-contamination, but we did identify one sample swap between two donors. We used featureCounts v1.5.0 (48) to count the number of uniquely mapping fragments overlapping GENCODE (49) basic annotation from Ensembl 79. We excluded short RNAs and pseudogenes from the analysis leaving 35,033 unique genes of which 19,796 were protein coding.

Furthermore, we only used 15,797 genes with mean expression in at least one of the conditions greater than 0.5 transcripts per million (TPM) (50) in all downstream analyses. We quantile-normalised the data and corrected for sample-specific GC content bias using the conditional quantile normalisation (cqn) (51) R package as recommended previously (52). To detect hidden confounders in gene expression, we applied PEER (53) to each condition separately allowing for at most 10 hidden factors. We found that the first 3-5 factors explained the most variation in the data and the others remained close to zero. Although we performed replicate macrophage differentiations and RNA-seq from four iPSC lines, for simplicity we decided to use only one of the replicates in downstream analyses. We further excluded samples from one donor (qaqx_1) from downstream analysis because they appeared as outliers in principal component analysis (PCA). The final dataset consisted of 336 RNA-seq samples from 84 donors.

#### Differential expression analysis

We included 15,797 genes whose mean expression in at least one of the conditions was greater than 0.5 TPM into our differential expression analysis. For each gene, we used likelihood ratio test implemented in DESeq2 (54) v1.10.0 (test = “LRT”) to test if a model that allowed different mean expression in each condition explained the data better than a null model assuming the same mean expression across conditions. Overall, 8758 genes with Benjamini-Hochberg FDR < 1% and fold change between naive and any one of the stimulated conditions greater than 2 were identified as differentially expressed.

To identify differentially expressed genes with specific expression patterns, we calculated mean quantile-normalised expression level in each condition and standardised the mean expression values across conditions to have zero mean and unit variance. Subsequently, we used c-means fuzzy clustering implemented in MFuzz v.2.28 (55) package with parameters ‘c = 9, m = 1.5, iter = 1000’ to assign the genes into 9 clusters. The number of clusters was chosen iteratively by trialing different numbers and observing which ones led to stable clustering results from independent runs. We ranked the genes in each cluster by their fold change between naive and highest expression conditions and used g:Profiler (56) R package with ‘max_set_size = 3000, ordered_query = TRUE, exclude_iea = TRUE’ options to identify pathways and Gene Ontology (GO) categories enriched in each cluster.

### ATAC-seq

#### Experimental procedures

We used an adapted version of the original ATAC-seq protocol (57). Approximately 150,000 cells were seeded into 1 well of a 6-well plate and treated identically to the RNA-seq samples. After stimulation, cells were washed once with ice-cold Dulbecco's phosphate buffered saline without calcium and magnesium and incubated for 12 minutes on ice in 500 µl freshly-made sucrose buffer (10mM Tris-Cl pH 7.5, 3 mM CaCl_2_, 2mM MgCl_2_, 0.32 M sucrose). After 12 minutes, 25 µl of 10% Triton-X-100 (final concentration 0.5%) was added and the cells were incubated for another 6 minutes to release the nuclei. The lysate was transferred to 1.5 mL microfuge tube, vortexed briefly, and centrifuged at 300 *g* for 8 minutes at 4°C. All traces of the sucrose lysis buffer were removed before immediately resuspending the nuclei pellet in 50 µL of Nextera tagmentation master mix (Illumina FC-121-1030), comprising 25 µL 2x Tagment DNA buffer, 20 µL nuclease-free water and 5 µL Tagment DNA Enzyme 1. The tagmentation reaction mixture was immediately transferred to a 1.5 mL low adherence microfuge tube and incubated at 37°C for 30 minutes. The tagmentation reaction was stopped by the addition of 250 µL Buffer PB (Qiagen) and PCR was performed as described in (23). The tagmented chromatin was then purified using the MinElute PCR purification kit (Qiagen 28004), according to the manufacturer's instructions, eluting in 10 µL of buffer EB (Qiagen). Finally, size selection was performed using agarose gel and SPRI beads (23). Five samples were pooled per lane and 75 bp paired end reads were sequenced on Illumina HiSeq 2000 using the V4 chemistry.

#### Read alignment

Illumina Nextera sequencing adapters were trimmed using skewer v0.1.127 (58) in paired end mode. Trimmed reads were aligned to GRCh38 human reference genome using bwa mem v0.7.12 (59). Reads mapping to the mitochondrial genome and alternative contigs were excluded from all downstream analysis. Picard 1.134 MarkDuplicates was used to remove duplicate fragments. We used verifyBamID (47) 1.1.2 to detect and correct potential sample swaps between individuals. Fragment coverage BigWig files were constructed using bedtools v2.17.0 (60).

#### Peak calling

We used MACS2 (61) v2.1.0 with ‘--nomodel --shift -25 --extsize 50 -q 0.01’ to identify open chromatin regions (peaks) that were enriched for transposase integration sites compared to the background at 1% FDR level. With these parameters we detected between 31,658 and 208,330 peaks per sample. We constructed consensus peak sets in each condition separately by pooling all of the peak calls from all of the samples. For each peak, we first counted the number of samples in which that peak was identified. We then calculated the union of all peaks that were detected in at least three samples. Finally, we pooled the consensus peaks from all four conditions to obtain the final set of 296,220 unique peaks that were used for all downstream analyses. We used featureCounts (48) v.1.5.0 to count the number of fragments overlapping consensus peak annotations and ASEReadCounter (62) from Genome Analysis Toolkit (GATK) to quantify allele-specific chromatin accessibility.

#### Sample quality control

We used the following criteria summarised in table S6 to assess the quality of ATAC-seq samples:

- *Assigned fragment count* - the total number of paired end fragments assigned to peaks by featureCounts.
- *Mitochondrial fraction* - fraction of total fragments aligned to the mitochondrial genome.
- *Assigned fraction* - fraction of non-mitochondrial reads assigned to consensus peaks. A measure of signal-to-noise ratio.
- *Duplicated fraction* - fraction of fragments that were marked as duplicates by Picard MarkDuplicates.
- *Peak count* - number of peaks called by MACS2.
- *Length ratio* - # of short fragments (< 150 nt) / # long fragments (> = 150 nt). This measures if the read length distribution has characteristic ATAC-seq profile with clearly visible mono-nucleosomal and di-nucleosomal peaks.

We used these criteria to exclude 5 samples from downstream analysis (table S6). One sample was excluded because of very low assigned fraction (∼10%) and peak count, two more were excluded because of extremely large length ratio (>7) and a fragment length distribution uncharacteristic for ATAC-seq library. The final two samples were excluded because they appeared to be outliers in the principal component analysis (PCA).

#### Differentially accessible regions

We used limma voom v3.26.3 (63) with TMM normalisation to identify 63,430 peaks that were more than 4-fold differentially accessible (FDR < 0.01) between naive and any one of the stimulated conditions. We only included high quality samples from 16 donors (64 samples) (table S6) in the analysis, because we noticed that limma voom was sensitive to additional noise in the lower quality samples. Subsequently, we quantile-normalised the peak accessibility data using cqn (51), calculated the mean accessibility of each peak in each condition and used Mfuzz v.2.28 (55) to cluster the peaks into seven distinct activity patterns.

#### Motif enrichment

We downloaded the CIS-BP (64) human TF motif database from the MEME website and used FIMO (65) to identify the occurrences of all TF motifs within the ATAC consensus peaks with FIMO threshold p-value < 1e-5. We also performed the same motif scan for 2kb promoter sequences upstream of 21,350 human genes (downloaded from the PWMEnrich (66) R package) and used this as the background set. We used Fisher’s exact test to identify motifs that occurred significantly more often in macrophage open chromatin regions compared to the background promoter sequences. Because the CIS-BP database contains many redundant motifs, we manually selected 21 representative motifs for downstream analysis corresponding to the major TFs important in macrophage biology: AP-1, IRF-family, ETS-family (PU.1, ELF1, FLI1), NF-κB, CEBPα, CEBPβ, ATF4, CTCF, STAT1, MAFB, MEF2A and USF1. We used Fisher’s exact test to identify motifs that were specifically enriched in each cluster of differentially accessible peaks compared to the background of all macrophage ATAC-seq peaks.

### ChIP-seq data analysis

Human macrophage PU.1 ChIP-seq data (27) (75 bp single-end reads) and C/EBPβ ChIP-seq data (67) (50 bp paired-end reads) were downloaded from GEO (accessions GSE66594 and GSE54975, respectively). Single-end dataset was aligned to the GRCh38 reference genome using bwa aln v0.7.12 and paired-end dataset was aligned to the same reference using bwa mem v0.7.12 with the -M flag set. For the paired-end dataset, only properly paired reads were used for downstream analysis. Duplicate reads were removed with Picard v1.134 MarkDuplicates with the ‘REMOVE_DUPLICATES = true’ parameter set. We used bedtools v2.17.0 (60) to construct genome wide read (single-end) or fragment (paired-end) coverage tracks in BigWig format. Finally, we called peaks using MACS2 v2.1.0 with ‘-q 0.01’ option.

### QTL mapping

#### Preparing genotype data

We obtained imputed genotypes for all of the samples from the HipSci (18) project. We used CrossMap v0.1.8 (68) to convert variant coordinates from GRCh37 reference genome to GRCh38. Subsequently, we filtered the VCF file with bcftools v.1.2 to retain only bi-allelic variants (both SNPs and indels) with IMP2 score > 0.4 and minor allele frequency (MAF) > 0.05 in our 86 samples. The same VCF file was used for all subsequent analyses. The genotype data for 52 managed access lines is available from the European Genome-phenome Archive (EGA) (EGAD00010000773), the data for the remaining 34 open access lines is deposited in the European Nucleotide Archive (ENA) (ERP013161). The VCF file was imported into R using the SNPRelate (69) R package.

#### Quantifying allele-specific expression and chromatin accessibility

We used ASEReadCounter (62) from the Genome Analysis ToolKit (GATK) to count the number of allele-specific fragments overlapping each variant in the RNA-seq and ATAC-seq datasets.

We used the following flags with ASEReadCounter: ‘-U ALLOW_N_CIGAR_READS -dt NONE -minMappingQuality 10 -rf MateSameStrand’. We removed indels from the VCF file prior to quantifying allele-specific expression because they are not supported by the RASQUAL model.

#### Detecting QTLs using RASQUAL

We wrote a collection of python scripts and a rasqualTools R package to simplify running RASQUAL on large number of samples and work with large RASQUAL output files. This software is available on GitHub (https://github.com/kauralasoo/rasqual). We used the vcfAddASE.py script to add allele-specific counts calculated in the previous step into the VCF file. We ran RASQUAL (23) independently for each experimental condition using sex and first two PEER factors as covariates (sex and first 3 PCs for caQTLs). In contrast to standard linear model, covariates seemed to have only a minor effect on the number of QTLs detected by RASQUAL. We only included variants that were either in the gene body or within +/-500kb from the gene (+/-50kb from the peak). We specified ‘--imputation-quality > 0.7’. As a result, variants with imputation quality of < 0.7 were used as feature SNPs in allele-specific analysis but were not considered as possible causal variants. We also used RASQUAL's GC correction option to correct for sample-specific GC bias in the feature-level read count data. To correct for multiple testing, we picked one minimal p-value per feature, used eigenMT (70) to estimate the number of independent tests performed in the cis-region of each feature and then performed Bonferroni correction to obtain the corrected p-value. We also ran RASQUAL once with the ‘--random-permutation’ option to obtain empirical null p-values from data with permuted sample labels. We performed the same eigenMT multiple testing procedure on the permuted p-values and compared the true association p-values to the empirical null distribution to identify QTLs with FDR < 10%.

#### Detecting QTLs using a linear model

We used linear regression implemented in the FastQTL (71) software to map cis-QTLs in each experimental condition. We used the ‘–permute 100 10000’ option to obtain permutation p-values for each association. The size of the cis windows was set to +/-500kb around each gene and +/-50kb around each ATAC-seq peak. Prior to to QTL mapping, the read count data was quantile normalised using the cqn package with GC-content of the feature (gene or peak) included as a covariate. For eQTL analysis, we used sex and the first six PEER factors as covariates in the model. For caQTL analysis we used sex and the first three principal components (PCs) as covariates in the model. Although FastQTL reported feature-level permutation p-values, obtaining those was computationally not feasible for RASQUAL. Therefore, to be able to faithfully compare the number of QTLs detected by FastQTL and RASQUAL, we decided to apply exactly the same multiple testing correction procedure (eigenMT + single permutation of sample labels) to both methods. We further restricted the comparison to features that were tested by both methods. This affected a small number of genes that were tested by FastQTL but filtered out by RASQUAL, because the raw read count was exactly zero in all samples.

#### Detecting response eQTLs

In each condition, we first identified all genes and corresponding lead variants that displayed significant association at 10% FDR level from RASQUAL. For each gene, we only kept independent lead variants (R^2^ < 0.8). Finally, we used all independent pairs of genes and corresponding lead variants to test if the eQTL effect size was significantly different between conditions. This was equivalent to testing the significance of the interaction term between condition and lead eQTL variant for each gene. Furthermore, to take advantage of the fact that gene expression was profiled in the same 84 lines in the four conditions, we also included the cell line as a random effect and fitted a linear mixed model using the lme4 (72) package.

Specifically, for each gene and lead variant pair we compared the following two models:

> H_0_: expression ∼ genotype + condition + covariates + (1|cell_line)
>
> H_1_: expression ∼ genotype + condition + genotype:condition + covariates + (1|cell_line)

where (1|cell_line) denotes the cell line specific random effect. We then calculated empirical p-values for the interaction test by permuting the conditions within each individual line 1,000 times. Subsequently, we used Benjamini-Hochberg FDR correction on the permutation p-values to identify 1,950 significant interactions at 10% FDR level. We used the same normalised data and covariates for interaction testing that were previously used for eQTL mapping in each condition separately.

#### Detecting response caQTLs

The procedure to identify response caQTLs was almost identical to the one used to detect response eQTLs above. However, instead of a linear mixed model we decided to use standard linear model without the random effect for cell line because not all lines were measured in all conditions. Furthermore, we found that our strategy to permute conditions within individual lines was not reliable when the number of measured conditions was not the same for each individual. Therefore, we decided to apply Benjamini-Hochberg FDR correction to nominal p-values from the linear model and identify significant interactions at 10% FDR level. With this approach we identified significant interactions for 6,591 caQTL regions.

#### Filtering and clustering QTLs based on effect size

Next, we focussed on all significant response eQTLs and response caQTLs that were detected with the interaction test above. We extracted the RASQUAL QTL effect size estimates π for each feature-variant pair in each condition and converted them to log_2_ fold changes between the two homozygotes using the formula log_2_FC = -log_2_(π/(1-π)). For a QTL to be considered condition specific we required the absolute log_2_FC in the naive condition to be less than 0.59 (1.5-fold) and the absolute difference in log_2_FC between naive and any one of the stimulated conditions to be greater than 0.59 (∼1.5 fold). We further required the absolute log_2_FC to be greater than 0.59 in at least one condition. To obtain relative log_2_FC, we divided the log_2_FC values in each condition by the maximal log_2_FC value observed across conditions. This scaling was necessary to make QTLs with different absolute effect size comparable to each other.

Finally, we used k-means clustering to identify six groups of QTLs that had similar activity patterns across conditions.

#### Sharing of QTL lead variants between conditions

To quantify how often QTL lead variants were shared between two conditions (e.g. A and B), we first identified all features (genes or peaks) with the nominal p-value of the lead variant < 1 × 10^−6^ in condition A. Next, for each feature, we took the lead variant in condition B and counted how often were the two lead variants in high LD with each other (R^2^ > 0.8). We did not impose any p-value threshold for the lead variant in condition B, reasoning that if there is no association in condition B then the lead variant is picked randomly and is therefore unlikely to be in high LD with the lead variant in condition A.

#### Identifying master and dependent regions

For each caQTL region, we defined its credible set of causal variants as those with R^2^ > 0.8 to the lead variant. We then classified the focal caQTL region as a master region (i - Fig. 3B), if the credible set overlapped the region itself, suggesting that the caQTL is directly caused by a variant within the region disrupting transcription factor binding. Alternatively, if the credible set overlapped some other regulated region but not the focal region, then we classified it as a dependent region (ii - Fig. 3B). We also excluded ambiguous cases where the credible set overlapped either multiple regulated regions (iii - Fig 3B) or it did not overlap any regulated regions (iv-v - Fig. 3B).

#### Motif disruption analysis

We limited motif disruption analysis to caQTL regions that did not contain associated indels and had < = 3 overlapping single nucleotide polymorphisms (SNPs) in them. For each SNP and peak pair we focussed on the sequence +/-25 bp from the SNP. We constructed both reference and alternative versions of the sequence and used TFBSTools v1.10.4 (73) to calculate the relative binding scores for both alleles (expressed as percentage from 0-100%). We considered the variant to be motif disrupting if the difference in relative binding score between the two alleles was > 3 percentage points. We also required the relative binding score for at least one of the alleles to be > = 85% of the theoretical maximum. This filter was necessary to exclude potential motif disruption events in very weak motif matches that were not likely to correspond to binding *in vivo* and is similar to the default thresholds recommended by TFBSTools. We used Fisher’s exact test to identify motifs that were significantly more often disrupted in one of the six condition-specific caQTL clusters compared to all caQTLs. For condition-specific caQTLs we further limited the analysis to putative master caQTL regions, because they were more likely to harbour the causal caQTL variant. We did not use that filter for caQTLs regulating putative primed enhancers, because the number of primed enhancers was much smaller.

#### Identifying condition-specific dependent peaks

To identify condition-specific dependent peaks, we tested if the effect size of the caQTL changed differently for master and dependent peaks (2,023 unique pairs) between two of conditions. This was equivalent to testing the significance of a three-way interactions between genotype, peak (master or dependent) and condition. We implemented this as the comparison of two standard linear models in R:

H0: accessibility ∼ genotype + peak + condition + peak*condition + genotype*peak + genotype*condition + covariates

H1: accessibility ∼ genotype + peak + condition + peak*condition + genotype*peak + genotype*condition + genotype*condition*peak + covariates

Similarly to condition-specific caQTL analysis, we used the first three principal components calculated separately for each condition as covariates in the model. For each master and dependent peak pair we picked the minimal p-value from three tests (naive vs each simulated condition) and used Bonferroni correction to correct for multiple testing. We then applied the Benjamini-Hochberg FDR correction to the Bonferroni-corrected p-values to identify all master-dependent peak pairs that showed significant interaction at 10% FDR. We used the log_2_FC from RASQUAL as the measure of caQTL effect size. To identify true condition-specific dependent peaks, we further filtered the results by requiring the absolute log_2_FC of the master peak to be >0.59 (1.5-fold) in the naive condition and the change in the log_2_FC for the dependent peak between the naive and stimulated condition to be > 0.59. We also required the change in the log_2_FC for the master peak to be < 1.

### Linking response eQTLs to caQTLs

First, we grouped all response eQTLs into three groups according to the condition in which they had the maximal effect size (IFNɣ, *Salmonella* or IFNɣ + *Salmonella*). For each response eQTL, we then identified all caQTLs that were in high linkage disequilibrium (LD) with it in any of the four conditions (R^2^ > 0.8 between the lead variants). If there was more than one caQTL in high LD, we picked the one with the smallest association p-value to obtain at most one caQTL corresponding to each response eQTL. Next, to estimate the prevalence of enhancer priming, we asked how often was the corresponding caQTL present already in the naive condition. Since response eQTLs were required to have RASQUAL log_2_FC < 0.59 in the naive condition (see above), we used the same threshold to decide the the caQTL was present (log_2_FC > 0.59) or absent (log_2_FC < 0.59) in the naive condition. Since there are various reasons why this analysis might lead to false positives, we decided to quantify our false positive rate by performing a reverse analysis where we started with response caQTLs, identified corresponding eQTLs (R^2^ > 0.8) and asked how often was the eQTL present already in the naive condition (log_2_FC > 0.59).

### Overlap with genome-wide association studies

We obtained full summary statistics for ten immune-mediated disorders: inflammatory bowel disease (IBD) including ulcerative colitis (UC) and Crohn's disease (CD) (74), Alzheimer's disease (AD) (75), rheumatoid arthritis (RA) (76), systemic lupus erythematosus (SLE) (77), type 1 diabetes (T1D) (78), schizophrenia (SCZ) (79), multiple sclerosis (MS) (80), celiac disease (CEL) (81) and narcolepsy (NAR) (82). In addition, we used summary statistics from type 2 diabetes (T2D) (83) as a negative control for a trait that should not be specifically enriched in macrophages. Summary statistics for T1D, CEL, IBD, RA, AD, MS and SLE were downloaded in 2015. SCZ, T2D and NAR were downloaded in 2016. T2D summary statistics were converted from GRCh36 to GRCh37 coordinates using the LiftOver tool, all of the other summary statistics already used GRCh37 coordinates.

Summary statistics for Alzheimer's disease were downloaded from International Genomics of Alzheimer's Project (IGAP). IGAP is a large two-stage study based upon genome-wide association studies (GWAS) on individuals of European ancestry. In stage 1, IGAP used genotyped and imputed data on 7,055,881 single nucleotide polymorphisms (SNPs) to meta-analyse four previously-published GWAS datasets consisting of 17,008 Alzheimer's disease cases and 37,154 controls (The European Alzheimer's disease Initiative – EADI the Alzheimer Disease Genetics Consortium – ADGC The Cohorts for Heart and Aging Research in Genomic Epidemiology consortium – CHARGE The Genetic and Environmental Risk in AD consortium – GERAD). In stage 2, 11,632 SNPs were genotyped and tested for association in an independent set of 8,572 Alzheimer's disease cases and 11,312 controls. Finally, a meta-analysis was performed combining results from stages 1 & 2.

#### Enrichment analysis

In each of the four conditions, we took all variants that were associated with either gene expression or chromatin accessibility with nominal p-value < 1e-5 and used that set of variants as a custom annotation track. We then used GARFIELD (84) to assess if all of these variants were collectivity enriched for GWAS hits for the ten traits described above. We excluded the MHC region (GRCh37: 6:20,000,000-40,000,000) from the GWAS summary statistics prior to enrichment testing, because this region was found to significantly inflate estimates of fold enrichment. In the main text, we reported fold enrichment at 1e-5 GWAS significance threshold.

#### Colocalisation analysis

We used coloc v2.3-1 (28) to test for colocalisation between molecular QTLs and GWAS hits. In the colocalisation analysis we used summary statistics from the linear model (rather than RASQUAL), because RASQUAL summary statistics could not be easily converted to approximate Bayes factors required by coloc. We ran coloc on a 400kb region centered on each lead eQTL and caQTL variant (200kb for the secondary eQTLs) that was less than 100kb away from at least one GWAS variant with nominal p-value < 1e-5. We then applied a set of filtering steps to identify a stringent set of eQTLs and caQTL that colocalised with GWAS hits. Similarly to what was done in (33), we first removed all cases where PP3 + PP4 < 0.8, to exclude loci where we were underpowered to detect colocalisation. We then required PP4/(PP3 + PP4) > 0.9 to only keep loci where coloc strongly prefered the model of a single shared causal variant driving both association signals over a model of two distinct causal variants. We excluded all colocalisation results from the MHC region (GRCh38: 6:28,510,120-33,480,577) because they were likely to be false positives due to complicated LD patterns in this region. We only kept results where the minimal GWAS p-value was < 10^−6^. Finally, we manually excluded 11 potential eQTL overlaps and 6 potential caQTL overlaps where on visual inspection the LD block exceeded the 400kb window that we used for colocalisation testing.

### Software used

Most figures were made in R using ggplot2 (85) and wiggleplotr (https://bioconductor.org/packages/wiggleplotr/) was used to produce RNA-seq and ATAC-seq read coverage plots. List of software used in this study is presented below.

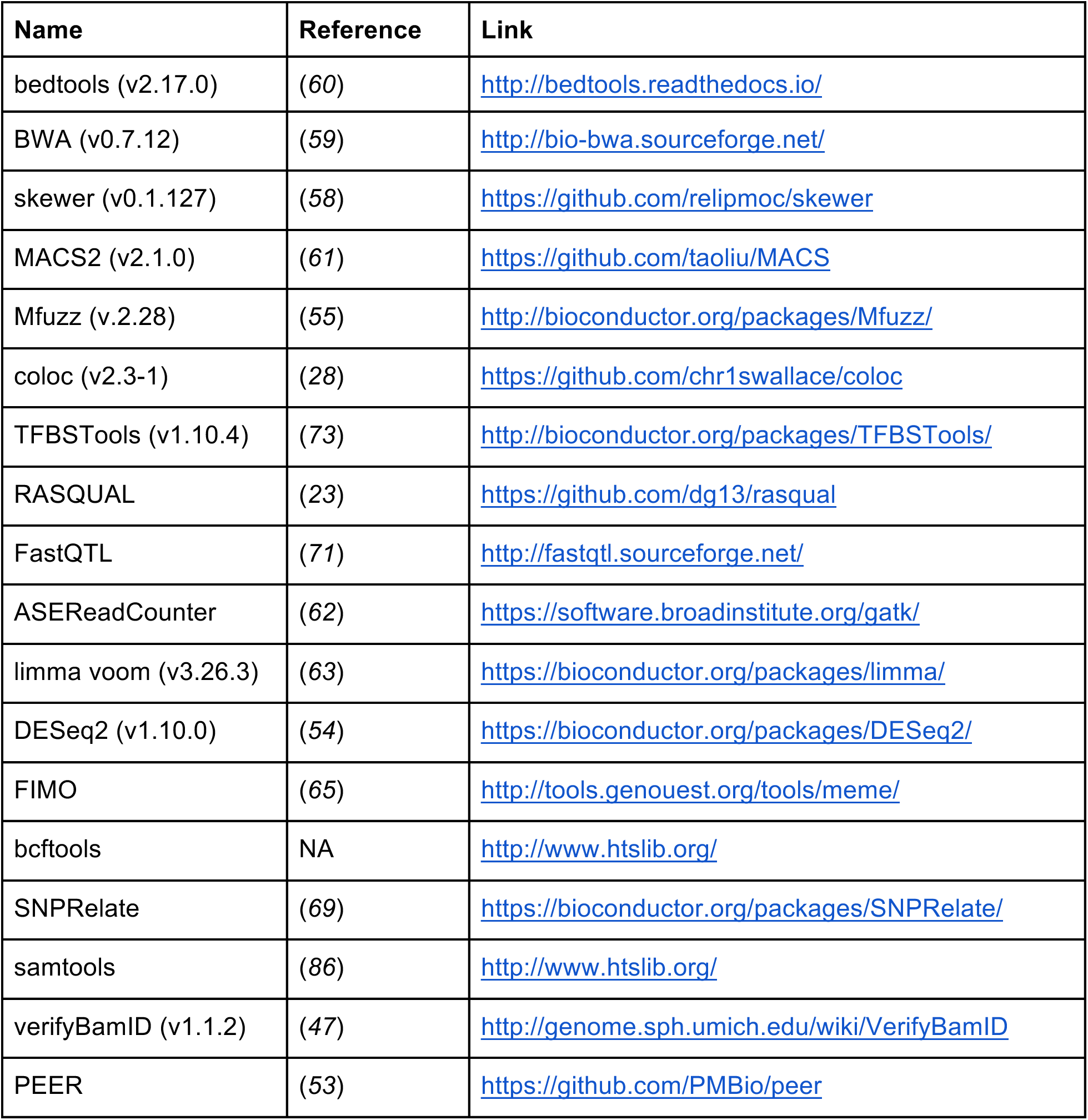

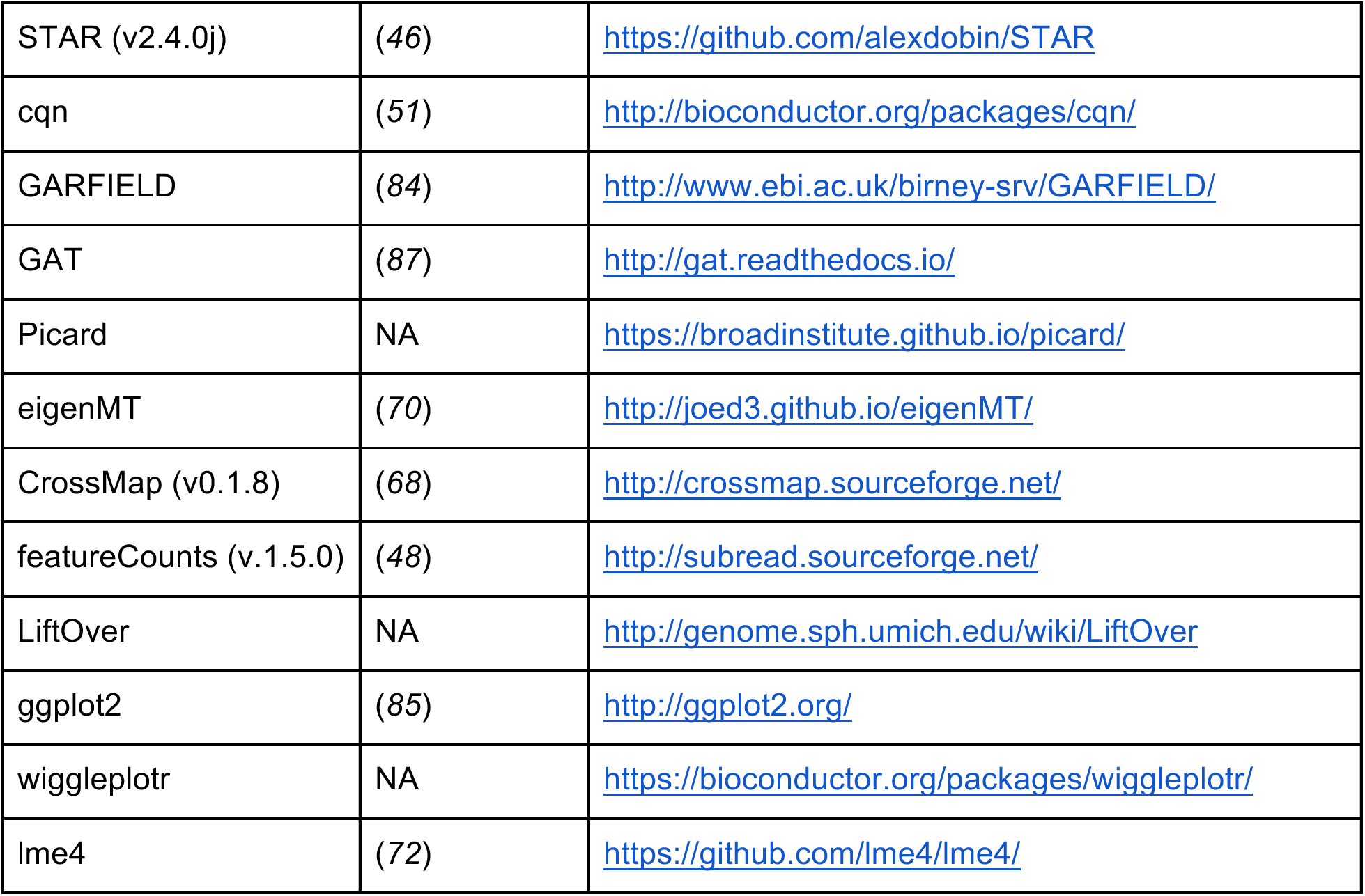

### Data availability

All of the experimental data has been deposited in the following repositories:

Data availability All of the experimental data has been deposited in the following repositories:
| Data | Managed access | Open access |
| --- | --- | --- |
| Imputed genotypes | EGAD00010000773 | ERP013161 |
| RNA-seq / ATAC-seq | EGAS00001002236 | ERP020977 |
| Flow cytometry | doi: 10.5281/zenodo.234214 |  |

## Supplementary Tables

Table S1: Metadata for all iPSC to macrophage differentiation attempts.

Table S2: GARFIELD fold enrichments and p-values for 10 different GWAS traits.

Table S3: List of eQTLs colocalised with GWAS associations.

Table S4: List of caQTLs colocalised with GWAS associations.

Table S5: Metadata for the RNA-seq samples.

Table S6: Metadata for the ATAC-seq samples.

## Supplementary Figures

**Figure S1:**
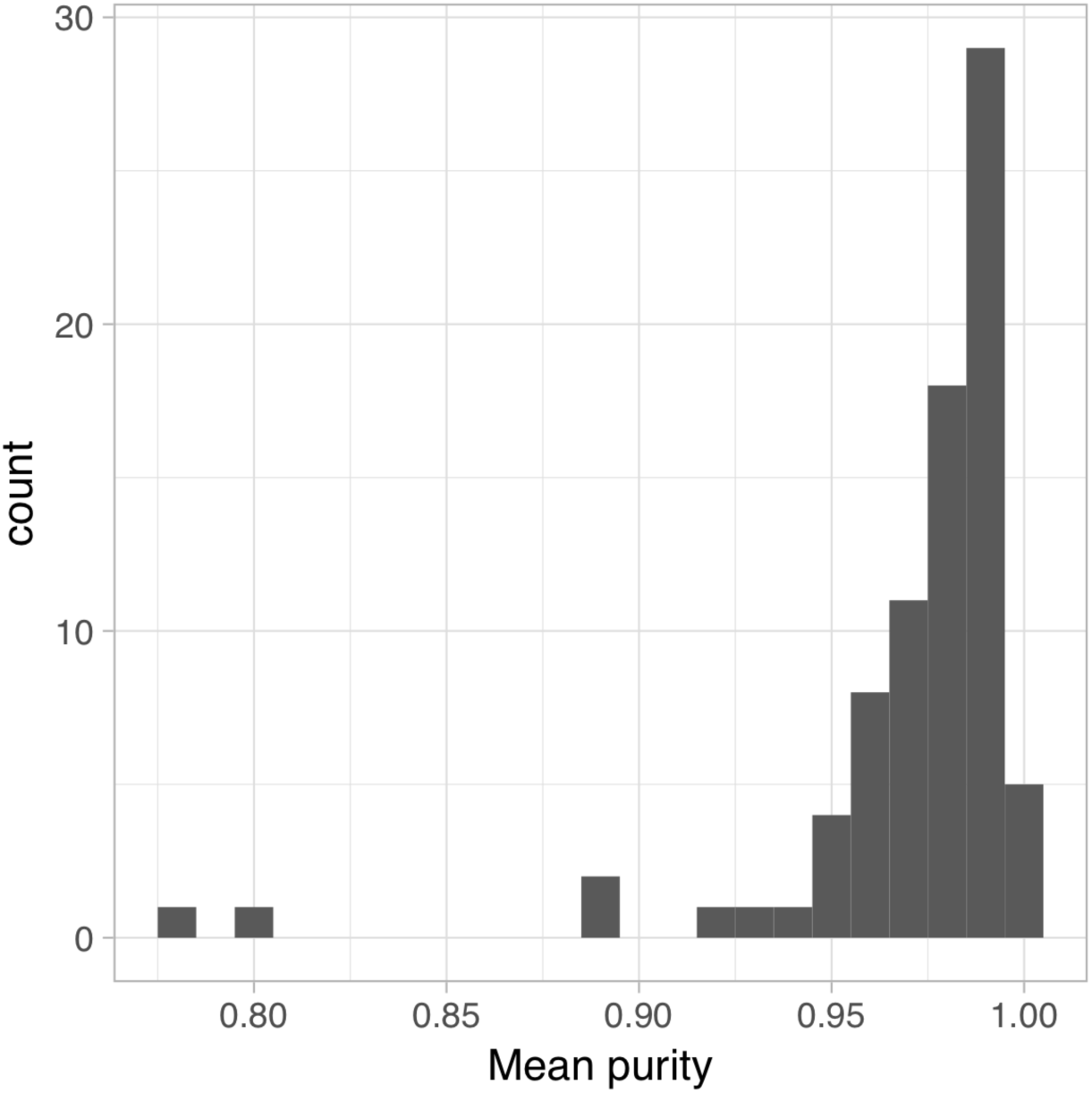
Histogram of the purity estimates of IPSC-derived macrophages. Macrophages were stained with antibodies for CD14, CD16 and CD206 and flow cytometry was used to estimate the proportion of cells stained positive for each of the three markers (see Methods). Since the purity estimates from all three markers were highly correlated, we decided to calculate their mean as an estimate of sample purity. The values for each sample are presented in table S1.

**Figure S2:**
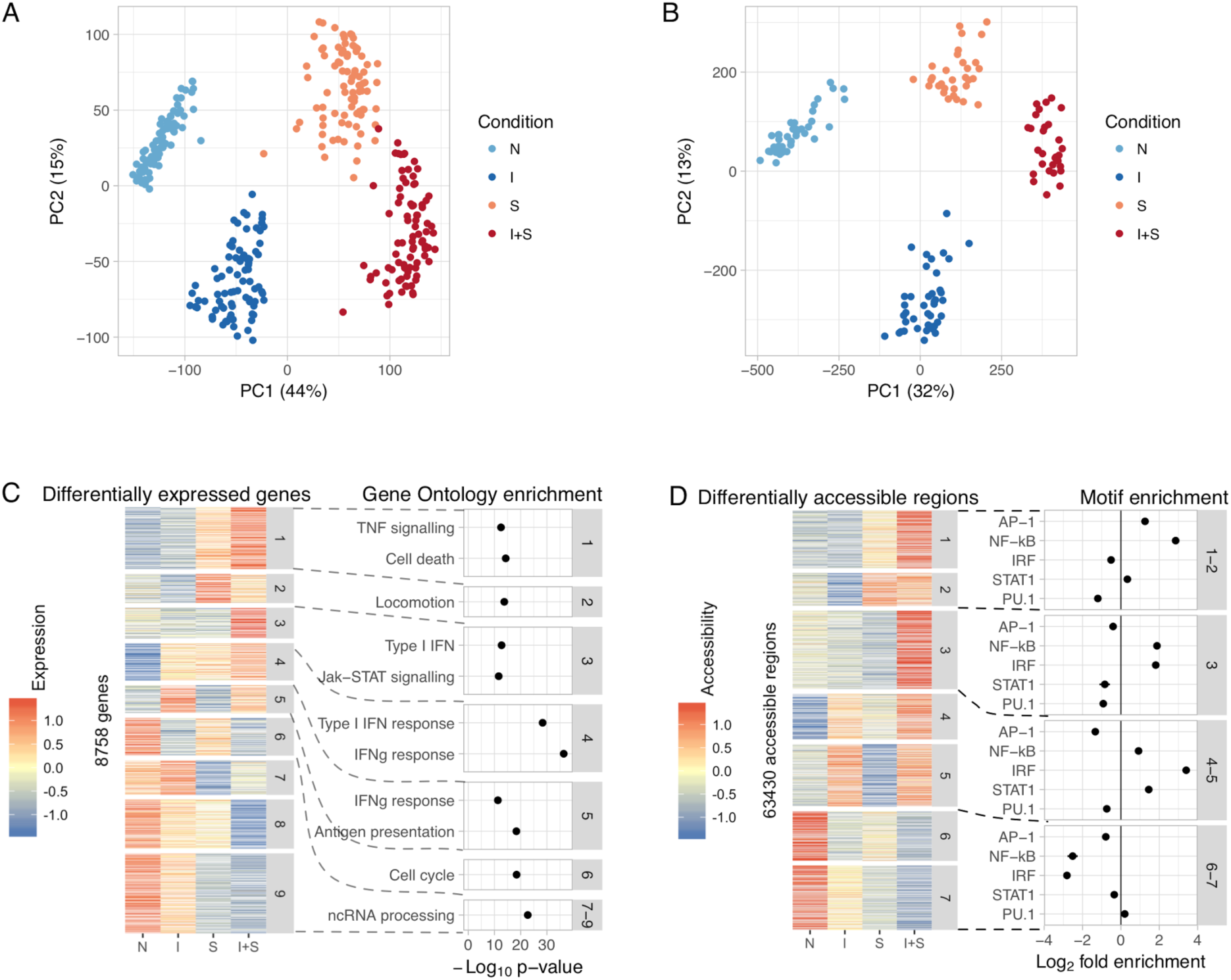
Differential gene expression and chromatin accessibility in macrophage immune response. **(A)** Principal component analysis of the gene expression data. **(B)** Principal component analysis of the chromatin accessibility data. **(C)** Left panel: 8,758 differentially expressed genes clustered into nine distinct expression patterns. Right panel: Selection of Gene Ontology terms enriched in each cluster. Only enrichments with p < 1×10^−8^ are shown in the figure. Differential gene expression patterns closely recapitulated known aspects of macrophage immune response. For example, genes upregulated by *Salmonella* (cluster 1) were enriched for tumor necrosis factor (TNF) signalling and cell death pathways whereas genes upregulated by IFNɣ (cluster 5) were enriched for IFNɣ response and antigen presentation pathways. **(D)** Left panel: heatmap of 63,350 differentially accessible regions clustered into seven distinct patterns. Right panel: enrichment of TF motifs in four groups of similar clusters relative to all open chromatin regions. Similarly to the gene expression data (panel A), open chromatin regions in clusters 1 and 2 that became accessible after *Salmonella* infection were specifically enriched for NF-κB and AP-1 motifs, two main TFs activated downstream of toll-like receptor 4 (TLR4) signalling (21). In contrast, clusters 4 and 5 showed increased accessibility after IFNɣ stimulation and were enriched for IRF and STAT1 motifs, consistent with the activation of STAT1 and IRF1 downstream of IFNɣ signalling (22). Solid lines represent 95% confidence intervals from Fisher’s exact test.

**Figure S3:**
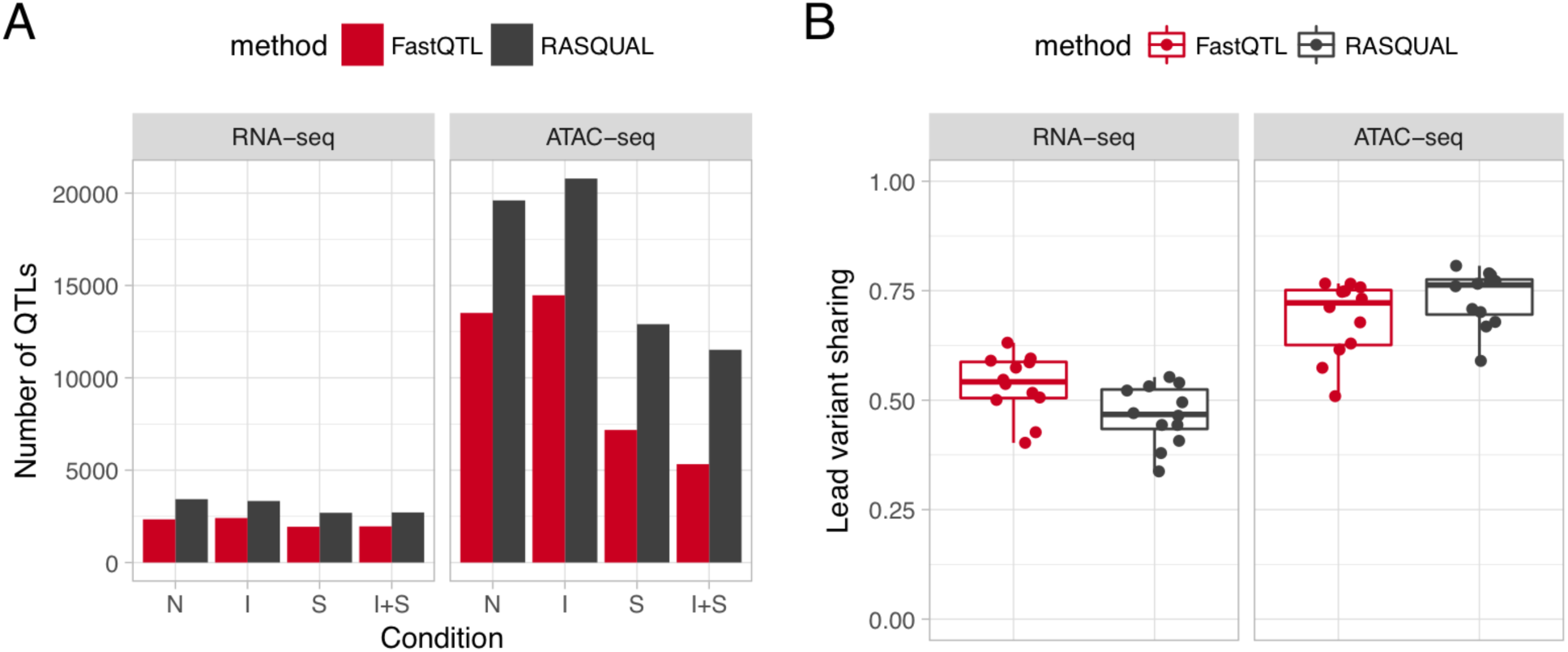
Genetic effects on gene expression and chromatin accessibility across conditions. **(A)** Number of genes and open chromatin regions for which we detected at least one significant QTL (FDR < 10%) using either allele-specific model (RASQUAL) or standard linear regression (FastQTL). The number of significant QTLs was counted in each condition separately. (**B**) Proportion of lead QTL variants shared between all pairs of conditions for both eQTLs and caQTLs.

**Figure S4:**
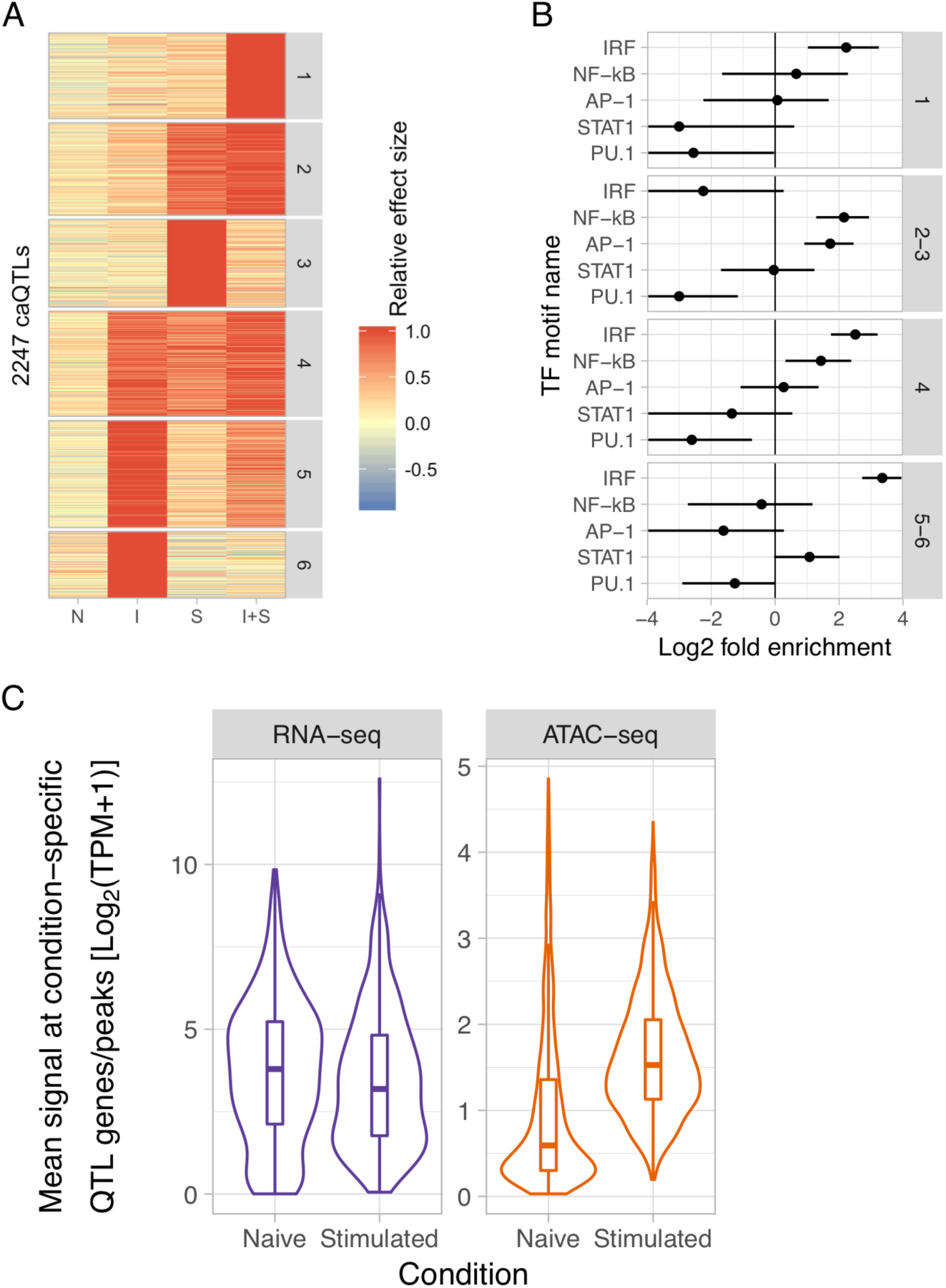
Characterising condition-specific caQTLs. **(A)** Condition-specific caQTLs clustered into six groups using k-means clustering. **(B)** *Salmonella*-specific caQTLs (clusters 2 and 3 from panel A) are enriched for disrupting NF-kB and AP-1 TF motifs whereas IFNɣ-specific caQTLs (clusters 5 and 6 from panel A) are enriched for disrupting the IRF motif. Solid lines represent 95% confidence intervals from Fisher’s exact test. **(C)** Distribution of mean gene expression (or chromatin accessibility) in the naive condition and in the condition where the condition-specific QTL had the largest effect size. Condition-specific caQTLs were inaccessible in the unstimulated cells and became accessible only in the condition in which the genetic effect appeared whereas mean expression of genes with condition-specific eQTLs did not differ between conditions. TPM, transcripts per million.

**Figure S5:**
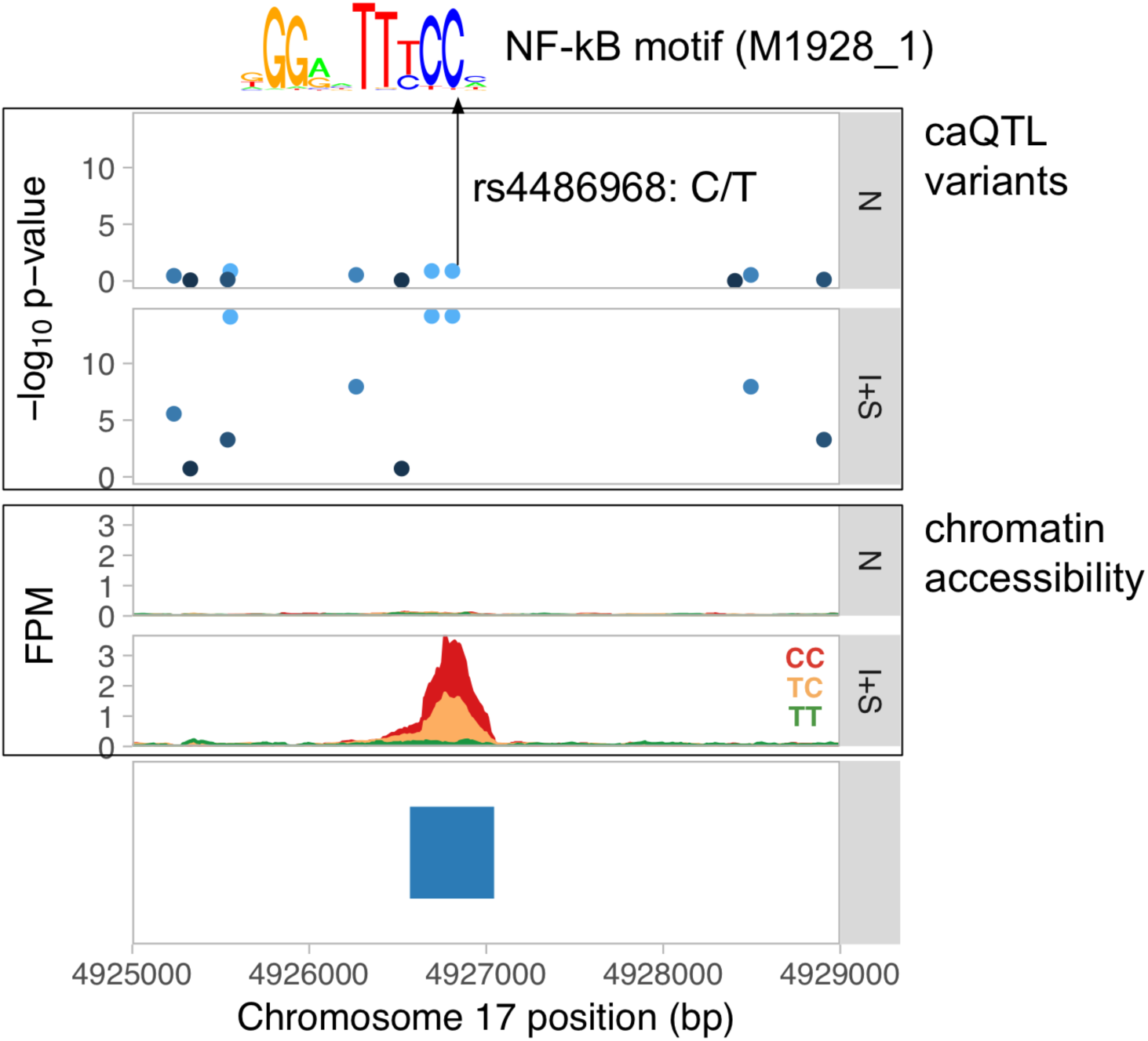
Fine mapping the causal variant for the caQTL upstream of GP1BA. The rs4486968 variant located within the accessible region is predicted to disrupt NF-κB binding motif (M1928_1 from CIS-BP (64)) by changing high affinity C (92.6% relative binding score) at position 10 to low affinity T (88.3% relative binding score).

**Figure S6:**
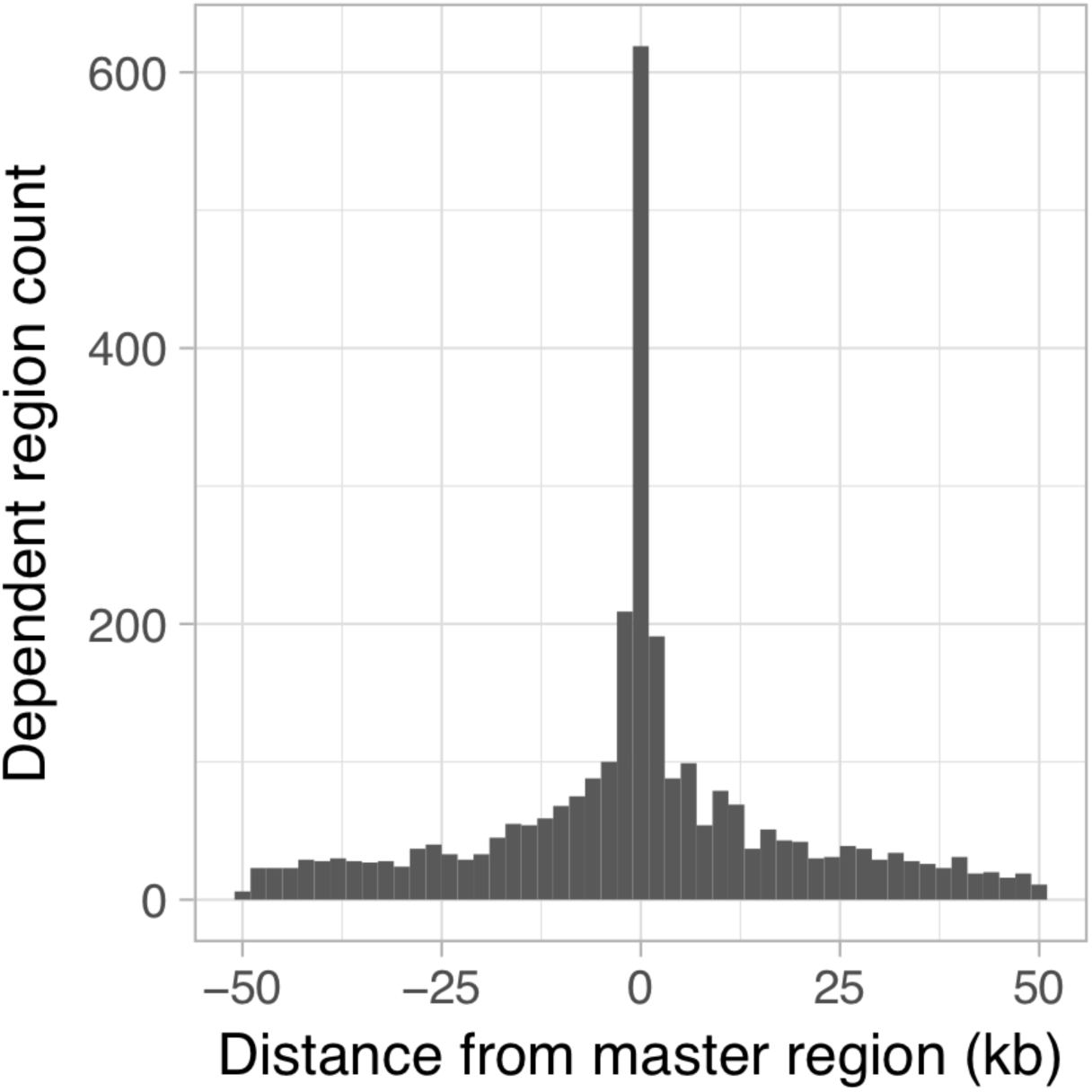
Histogram of the distances between master and dependent regions.

**Figure S7:**
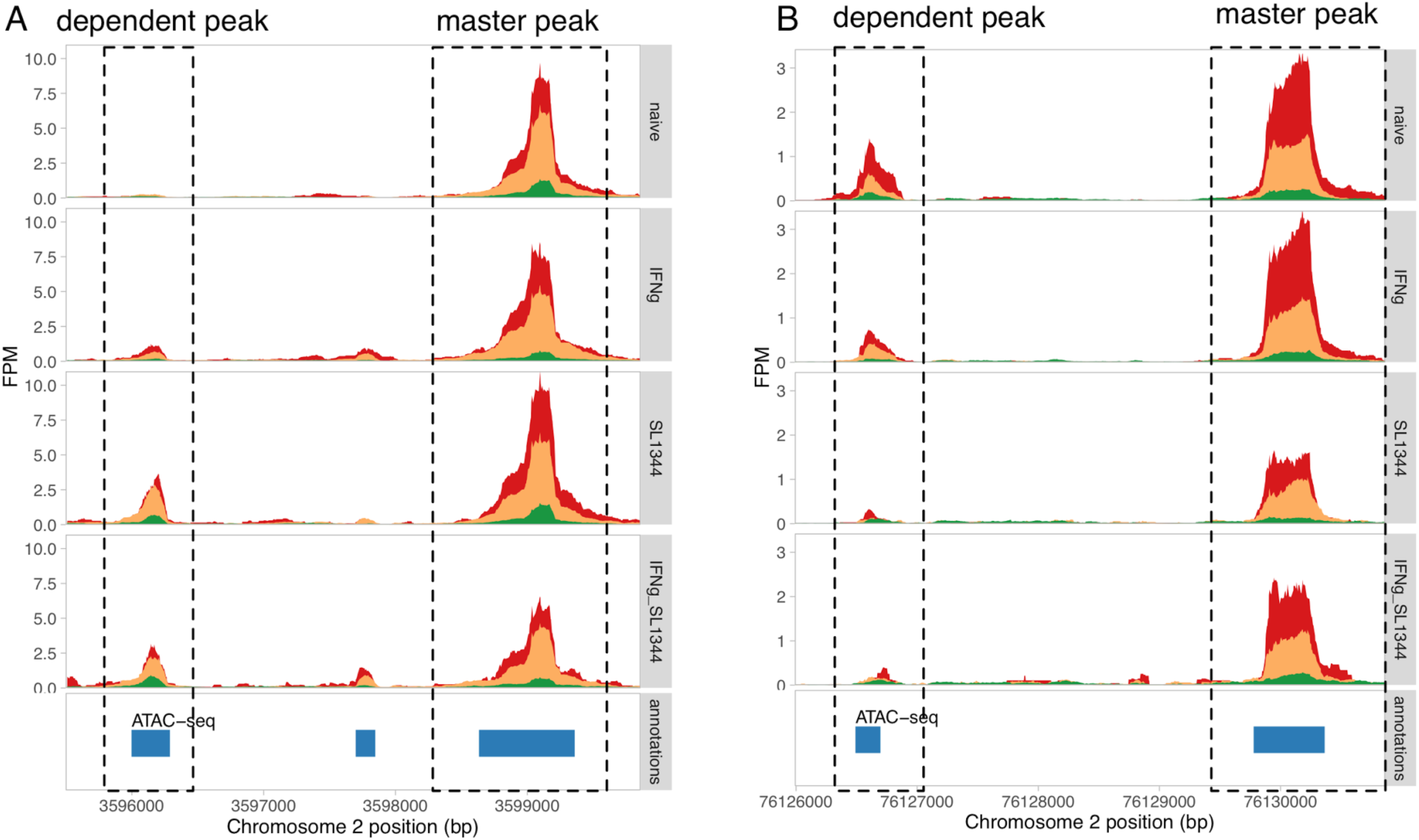
Two examples of condition-specific dependent peaks. (**A**) Dependent peak appears after *Salmonella* infection. (**B**) Dependent peak disappears after *Salmonella* infection.

**Figure S8:**
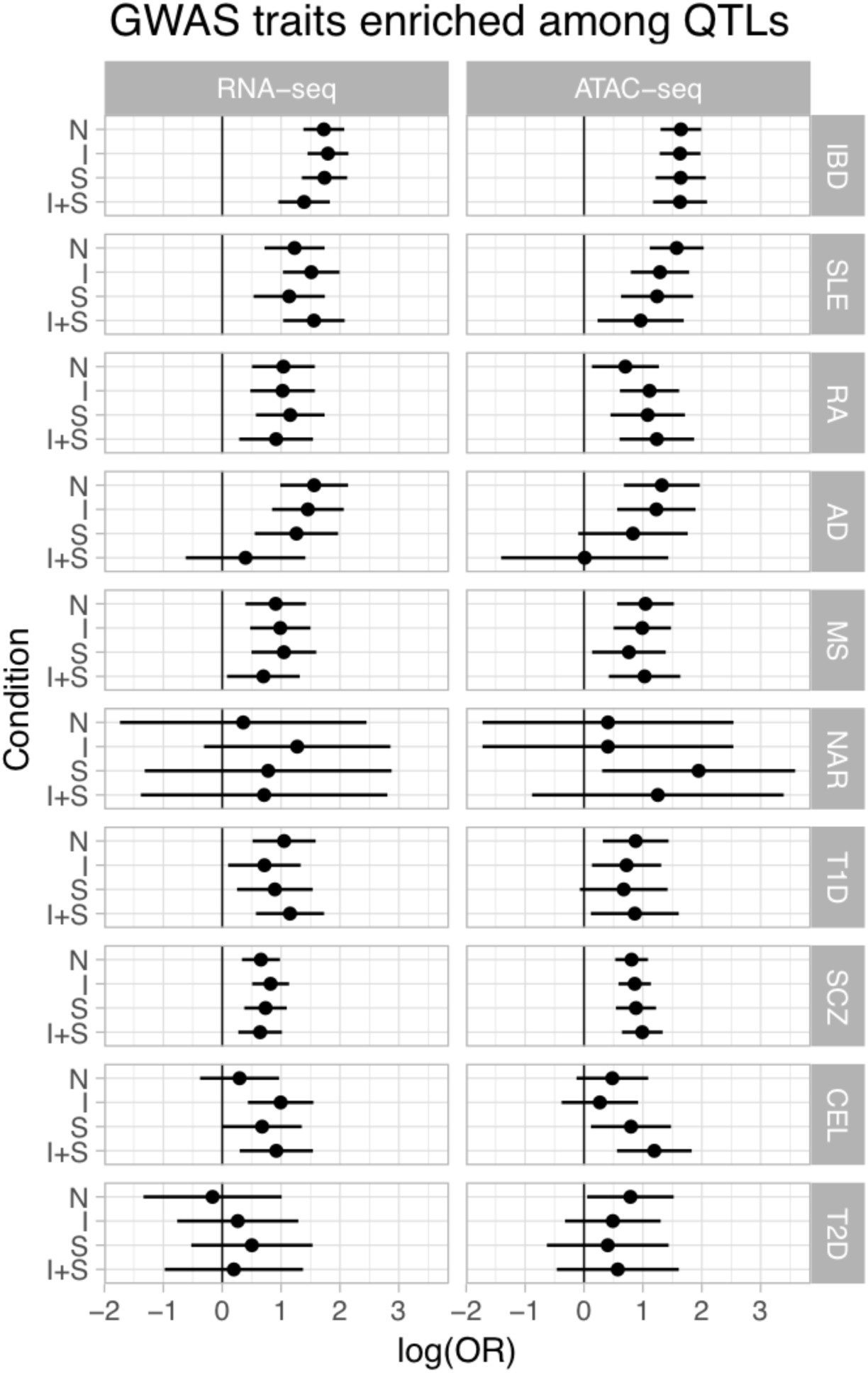
GARFIELD (84) enrichment of eQTLs and caQTLs among GWAS summary statistics from 10 complex diseases. The points represent the logarithm of enrichment odds ratios (OR) and error bars show the 95% confidence intervals. Disease acronyms: IBD, inflammatory bowel disease; RA, rheumatoid arthritis; SLE, systemic lupus erythematosus; AD, Alzheimer's disease; SCZ, schizophrenia; T2D, type 2 diabetes; MS, multiple sclerosis; NAR, narcolepsy; CEL, celiac disease; T1D, type 1 diabetes.

**Figure S9:**
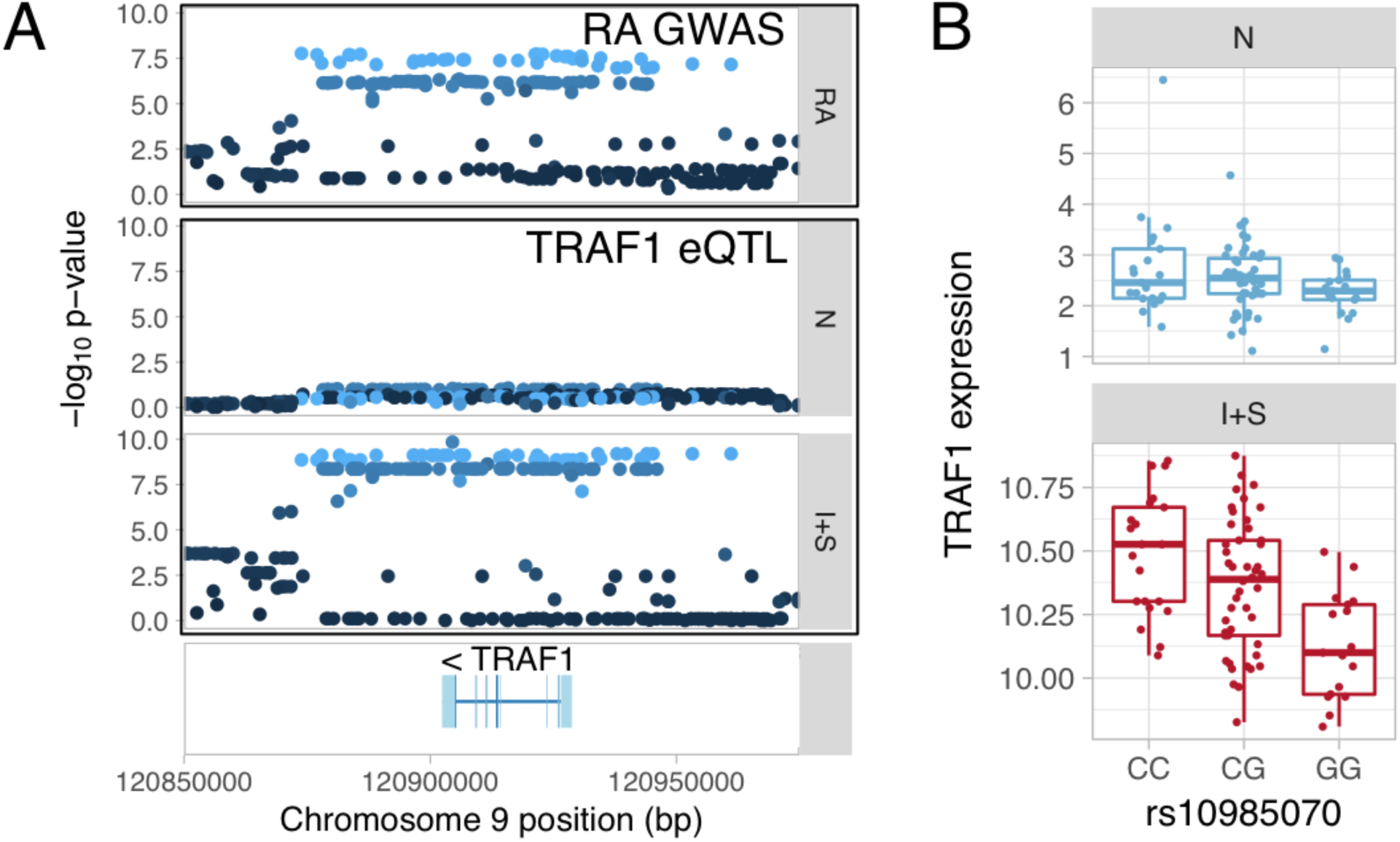
Colocalisation between a response eQTL and a GWAS hit. **(A)** IFNɣ + *Salmonella* specific eQTL for TRAF1 colocalised with a GWAS hit for rheumatoid arthritis (RA). (**B)** TRAF1 expression in naive and IFNɣ + *Salmonella* conditions stratified by the lead GWAS variant.

**Figure S10:**
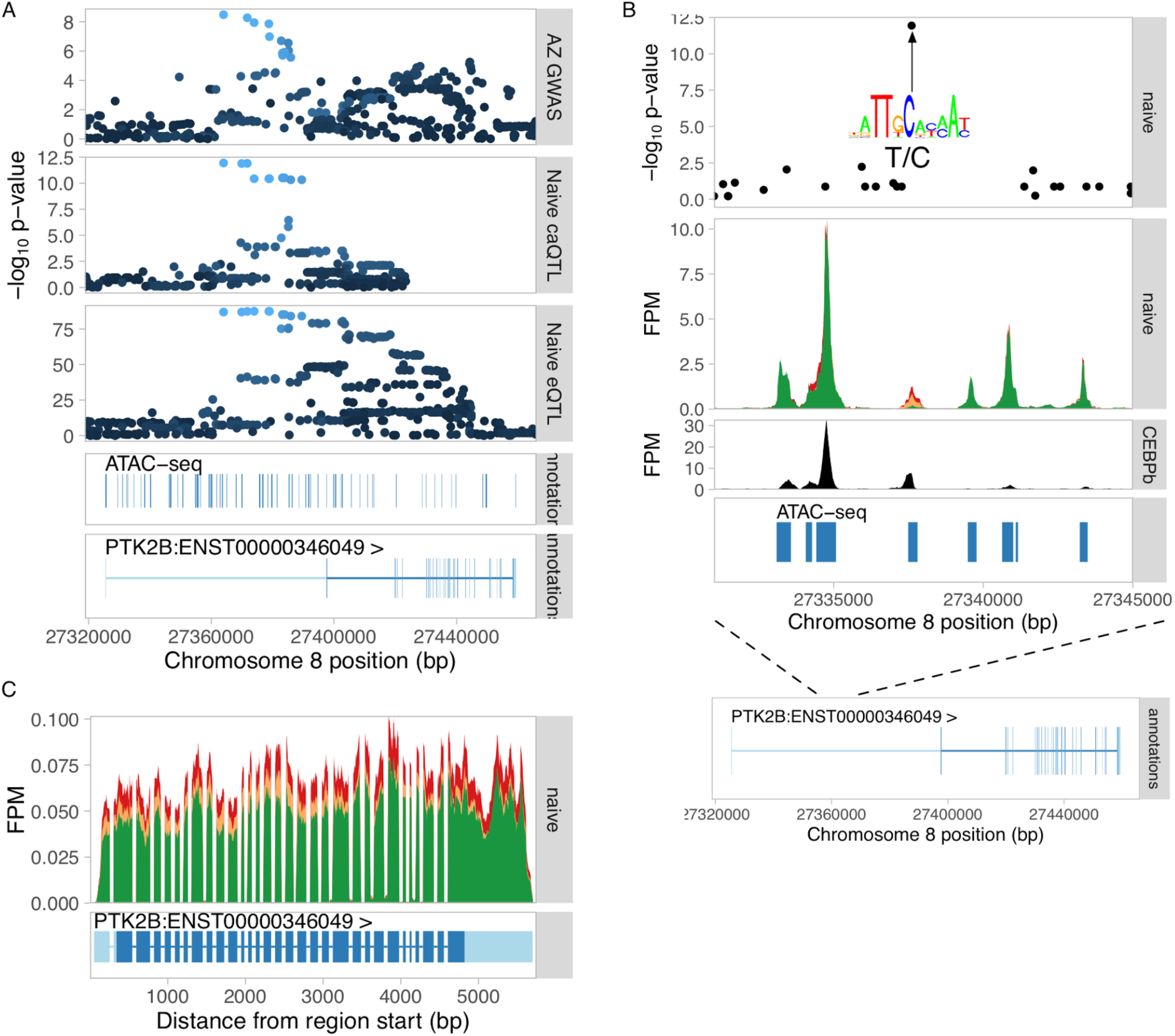
Dissecting the Alzheimer's disease causal variant at the PTK2B locus. (**A**) Manhattan plots for the Alzheimer's GWAS hit (top panel), colocalised caQTL (second panel) and colocalised eQTL for PTK2B gene (third panel). The bottom two tracks show all ATAC-seq peaks in the region as well was exons of the PTK2B gene. (**B**) ATAC-seq fragment coverage stratified by the lead caQTL and GWAS variant rs28834970. The lead variant was the only associated variant lying within the regulated caQTL peak, suggesting this is the most likely causal variant. The lead variant rs28834970 is T/C polymorphism and the alternative C allele is predicted to increase the relative binding score of the CEBPβ TF motif (M2268_1.02 in CIS-BP (64)) from 0.86 to 0.97. This is consistent with the increased chromatin accessibility at the C allele (middle panel) as well as increased expression of the PTK2B gene (panel C). Furthermore, the variant also overlaps experimental CEBPβ ChIP-seq peak in primary human macrophages (bottom panel) (67). Together, this evidence suggests that rs28834970 is the likely causal variant for Alzheimer's disease risk that influences PTK2B expression by disrupting CEBPβ motif in an enhancer in the first intron of the gene. (**C**) RNA-seq read coverage plot at the PTK2B gene stratified by the rs28834970 genotype.

## References

1. Y. Li et al., Inter-individual variability and genetic influences on cytokine responses to bacteria and fungi. Nat. Med. 22, 952–960 (2016).

2. L. B. Barreiro et al., Deciphering the genetic architecture of variation in the immune response to Mycobacterium tuberculosis infection. Proc. Natl. Acad. Sci. U. S. A. 109, 1204–1209 (2012).

3. P. Fairfax et al., Innate immune activity conditions the effect of regulatory variants upon monocyte gene expression. Science. 343, 1246949 (2014).

4. S. Kim et al., Characterizing the genetic basis of innate immune response in TLR4-activated human monocytes. Nat. Commun. 5, 5236 (2014).

5. M.N. Lee et al., Common genetic variants modulate pathogen-sensing responses in human dendritic cells. Science. 343, 1246980 (2014).

6. M. Çalışkan, S. W. Baker, Y. Gilad, C. Ober, Host genetic variation influences gene expression response to rhinovirus infection. PLoS Genet. 11, e1005111 (2015).

7. Y. Nédélec et al., Genetic Ancestry and Natural Selection Drive Population Differences in Immune Responses to Pathogens. Cell. 167, 657–669.e21 (2016).

8. K.M. de Lange et al., Genome-wide association study implicates immune activation of multiple integrin genes in inflammatory bowel disease. Nat. Genet. (2017), doi:10.1038/ng.3760.

9. S. Kim-Hellmuth et al., Genetic regulatory effects modified by immune activation contribute to autoimmune disease associations. bioRxiv (2017), p. 116376.

10. F. Degner et al., DNase I sensitivity QTLs are a major determinant of human expression variation. Nature. 482, 390–394 (2012).

11. F. Jin, Y. Li, B. Ren, R. Natarajan, PU.1 and C/EBP(alpha) synergistically program distinct response to NF-kappaB activation through establishing monocyte specific enhancers. Proc. Natl. Acad. Sci. U. S. A. 108, 5290–5295 (2011).

12. S. Heinz et al., Simple combinations of lineage-determining transcription factors prime cis-regulatory elements required for macrophage and B cell identities. Mol. Cell. 38, 576–589 (2010).

13. S. Heinz et al., Effect of natural genetic variation on enhancer selection and function. Nature. 503, 487–492 (2013).

14. A. Wang et al., Epigenetic priming of enhancers predicts developmental competence of hESC-derived endodermal lineage intermediates. Cell Stem Cell. 16, 386–399 (2015).

15. A. Chow, L. D. Jasenosky, A. E. Goldfeld, A Distal Locus Element Mediates IFN-γ Priming of Lipopolysaccharide-Stimulated TNF Gene Expression. Cell Rep. (2014), doi:10.1016/j.celrep.2014.11.011.

16. H.Y. Shin et al., Hierarchy within the mammary STAT5-driven Wap super-enhancer. Nat. Genet. 48, 904–911 (2016).

17. K. Alasoo et al., Transcriptional profiling of macrophages derived from monocytes and iPS cells identifies a conserved response to LPS and novel alternative transcription. Sci. Rep. 5, 12524 (2015).

18. H. Kilpinen et al., Common genetic variation drives molecular heterogeneity in human iPSCs. Nature (2017), doi:10.1038/nature22403.

19. X. Hu, L. B. Ivashkiv, Cross-regulation of signaling pathways by interferon-gamma: implications for immune responses and autoimmune diseases. Immunity. 31, 539–550 (2009).

20. Y. Qiao et al., Synergistic activation of inflammatory cytokine genes by interferon-γ-induced chromatin remodeling and toll-like receptor signaling. Immunity. 39, 454–469 (2013).

21. O. Takeuchi, S. Akira, Pattern recognition receptors and inflammation. Cell. 140, 805–820 (2010).

22. K. Schroder, P. J. Hertzog, T. Ravasi, D. A. Hume, Interferon-gamma: an overview of signals, mechanisms and functions. J. Leukoc. Biol. 75, 163–189 (2004).

23. N. Kumasaka, A. J. Knights, D. J. Gaffney, Fine-mapping cellular QTLs with RASQUAL and ATAC-seq. Nat. Genet. 48, 206–213 (2016).

24. F. Grubert et al., Genetic Control of Chromatin States in Humans Involves Local and Distal Chromosomal Interactions. Cell. 162, 1051–1065 (2015).

25. S.M. Waszak et al., Population Variation and Genetic Control of Modular Chromatin Architecture in Humans. Cell. 162, 1039–1050 (2015).

26. S. Cheng et al., Genetic determinants of chromatin accessibility and gene regulation in T cell activation across human individuals. bioRxiv, 090241 (2016).

27. S.V. Schmidt et al., The transcriptional regulator network of human inflammatory macrophages is defined by open chromatin. Cell Res. 26, 151–170 (2016).

28. Giambartolomei et al., Bayesian Test for Colocalisation between Pairs of Genetic Association Studies Using Summary Statistics. PLoS Genet. 10, e1004383 (2014).

29. Zhu et al., Integration of summary data from GWAS and eQTL studies predicts complex trait gene targets. Nat. Genet. 48, 481–487 (2016).

30. R. Jansen et al., Conditional eQTL analysis reveals allelic heterogeneity of gene expression. Hum. Mol. Genet. 26, 1444–1451 (2017).

31. M.T. Maurano et al., Systematic localization of common disease-associated variation in regulatory DNA. Science. 337, 1190–1195 (2012).

32. K. K.-H. Farh et al., Genetic and epigenetic fine mapping of causal autoimmune disease variants. Nature. 518, 337–343 (2015).

33. H. Guo et al., Integration of disease association and eQTL data using a Bayesian colocalisation approach highlights six candidate causal genes in immune-mediated diseases. Hum. Mol. Genet. 24, 3305–3313 (2015).

34. S. Chun et al., Limited statistical evidence for shared genetic effects of eQTLs and autoimmune-disease-associated loci in three major immune-cell types. Nat. Genet. 49, 600–605 (2017).

35. B.M. Javierre et al., Lineage-Specific Genome Architecture Links Enhancers and Non-coding Disease Variants to Target Gene Promoters. Cell. 167, 1369–1384.e19 (2016).

36. F. Jin et al., A high-resolution map of the three-dimensional chromatin interactome in human cells. Nature. 503, 290–294 (2013).

37. Y. Ghavi-Helm et al., Enhancer loops appear stable during development and are associated with paused polymerase. Nature. 512, 96–100 (2014).

38. C. Mullen et al., Master transcription factors determine cell-type-specific responses to TGF-β signaling. Cell. 147, 565–576 (2011).

39. Trompouki et al., Lineage regulators direct BMP and Wnt pathways to cell-specific programs during differentiation and regeneration. Cell. 147, 577–589 (2011).

40. R. Ostuni et al., Latent enhancers activated by stimulation in differentiated cells. Cell. 152, 157–171 (2013).

41. L. Magnani, J. Eeckhoute, M. Lupien, Pioneer factors: directing transcriptional regulators within the chromatin environment. Trends Genet. 27, 465–474 (2011).

42. R.I. Sherwood et al., Discovery of directional and nondirectional pioneer transcription factors by modeling DNase profile magnitude and shape. Nat. Biotechnol. 32, 171–178 (2014).

43. V. R. Ramirez-Carrozzi et al., A unifying model for the selective regulation of inducible transcription by CpG islands and nucleosome remodeling. Cell. 138, 114–128 (2009).

44. Bojcsuk, G. Nagy, B. L. Balint, Inducible super-enhancers are organized based on canonical signal-specific transcription factor binding elements. Nucleic Acids Res. 45, 3693–3706 (2017).

45. van Wilgenburg, C. Browne, J. Vowles, S. A. Cowley, Efficient, long term production of monocyte-derived macrophages from human pluripotent stem cells under partly-defined and fully-defined conditions. PLoS One. 8, e71098 (2013).

46. Dobin et al., STAR: ultrafast universal RNA-seq aligner. Bioinformatics. 29, 15–21 (2013).

47. G. Jun et al., Detecting and estimating contamination of human DNA samples in sequencing and array-based genotype data. Am. J. Hum. Genet. 91, 839–848 (2012).

48. Y. Liao, G. K. Smyth, W. Shi, featureCounts: an efficient general purpose program for assigning sequence reads to genomic features. Bioinformatics. 30, 923–930 (2014).

49. J. Harrow et al., GENCODE: the reference human genome annotation for The ENCODE Project. Genome Res. 22, 1760–1774 (2012).

50. P. Wagner, K. Kin, V. J. Lynch, Measurement of mRNA abundance using RNA-seq data: RPKM measure is inconsistent among samples. Theory Biosci. 131, 281–285 (2012).

51. K. D. Hansen, R. A. Irizarry, Z. Wu, Removing technical variability in RNA-seq data using conditional quantile normalization. Biostatistics. 13, 204–216 (2012).

52. S.E. Ellis et al., RNA-Seq optimization with eQTL gold standards. BMC Genomics. 14, 892 (2013).

53. O. Stegle, L. Parts, M. Piipari, J. Winn, R. Durbin, Using probabilistic estimation of expression residuals (PEER) to obtain increased power and interpretability of gene expression analyses. Nat. Protoc. 7, 500–507 (2012).

54. M. I. Love, W. Huber, S. Anders, Moderated estimation of fold change and dispersion for RNA-seq data with DESeq2. Genome Biol. 15, 550 (2014).

55. L. Kumar, M. E Futschik, Mfuzz: a software package for soft clustering of microarray data. Bioinformation. 2, 5–7 (2007).

56. J. Reimand et al., g:Profiler—a web server for functional interpretation of gene lists (2016 update). Nucleic Acids Res. 44, W83–W89 (2016).

57. J. D. Buenrostro, P. G. Giresi, L. C. Zaba, H. Y. Chang, W. J. Greenleaf, Transposition of native chromatin for fast and sensitive epigenomic profiling of open chromatin, DNA-binding proteins and nucleosome position. Nat. Methods. 10, 1213–1218 (2013).

58. Jiang, R. Lei, S.-W. Ding, S. Zhu, Skewer: a fast and accurate adapter trimmer for next-generation sequencing paired-end reads. BMC Bioinformatics. 15, 182 (2014).

59. Li, Aligning sequence reads, clone sequences and assembly contigs with BWA-MEM. arXiv [q-bio.GN] (2013) (available at http://arxiv.org/abs/1303.3997).

60. R. Quinlan, I. M. Hall, BEDTools: a flexible suite of utilities for comparing genomic features. Bioinformatics. 26, 841–842 (2010).

61. Y. Zhang et al., Model-based analysis of ChIP-Seq (MACS). Genome Biol. 9, R137 (2008).

62. S. Castel, A. Levy-Moonshine, P. Mohammadi, E. Banks, T. Lappalainen, Tools and best practices for data processing in allelic expression analysis. Genome Biol. 16, 195 (2015).

63. W. Law, Y. Chen, W. Shi, G. K. Smyth, voom: Precision weights unlock linear model analysis tools for RNA-seq read counts. Genome Biol. 15, R29 (2014).

64. M.T. Weirauch et al., Determination and Inference of Eukaryotic Transcription Factor Sequence Specificity. Cell. 158, 1431–1443 (2014).

65. C. E. Grant, T. L. Bailey, W. S. Noble, FIMO: scanning for occurrences of a given motif. Bioinformatics. 27, 1017–1018 (2011).

66. R. Stojnic, D. Diez, PWMEnrich: PWM enrichment analysis. R package version 4.8.2. *Bioconductor* (2015), (available at http://bioconductor.org/packages/release/bioc/html/PWMEnrich.html).

67. M.E. Reschen et al., Lipid-induced epigenomic changes in human macrophages identify a coronary artery disease-associated variant that regulates PPAP2B Expression through Altered C/EBP-beta binding. PLoS Genet. 11, e1005061 (2015).

68. H. Zhao et al., CrossMap: a versatile tool for coordinate conversion between genome assemblies. Bioinformatics. 30, 1006–1007 (2014).

69. X. Zheng et al., A high-performance computing toolset for relatedness and principal component analysis of SNP data. Bioinformatics. 28, 3326–3328 (2012).

70. J.R. Davis et al., An Efficient Multiple-Testing Adjustment for eQTL Studies that Accounts for Linkage Disequilibrium between Variants. Am. J. Hum. Genet. 98, 216–224 (2016).

71. Ongen, A. Buil, A. A. Brown, E. T. Dermitzakis, O. Delaneau, Fast and efficient QTL mapper for thousands of molecular phenotypes. Bioinformatics. 32, 1479–1485 (2016).

72. D. Bates, M. Mächler, B. Bolker, S. Walker, Fitting Linear Mixed-Effects Models Using lme4. J. Stat. Softw. 67, 1–48 (2015).

73. G. Tan, B. Lenhard, TFBSTools: an R/bioconductor package for transcription factor binding site analysis. Bioinformatics. 32, 1555–1556 (2016).

74. Z. Liu et al., Association analyses identify 38 susceptibility loci for inflammatory bowel disease and highlight shared genetic risk across populations. Nat. Genet. 47, 979–986 (2015).

75. C. Lambert et al., Meta-analysis of 74,046 individuals identifies 11 new susceptibility loci for Alzheimer's disease. Nat. Genet. 45, 1452–1458 (2013).

76. Y. Okada et al., Genetics of rheumatoid arthritis contributes to biology and drug discovery. Nature. 506, 376–381 (2014).

77. J. Bentham et al., Genetic association analyses implicate aberrant regulation of innate and adaptive immunity genes in the pathogenesis of systemic lupus erythematosus. Nat. Genet. 47, 1457–1464 (2015).

78. S. Onengut-Gumuscu et al., Fine mapping of type 1 diabetes susceptibility loci and evidence for colocalization of causal variants with lymphoid gene enhancers. Nat. Genet. 47, 381–386 (2015).

79. Schizophrenia Working Group of the Psychiatric Genomics Consortium, Biological insights from 108 schizophrenia-associated genetic loci. Nature. 511, 421–427 (2014).

80. International Multiple Sclerosis Genetics Consortium (IMSGC) et al., Analysis of immune related loci identifies 48 new susceptibility variants for multiple sclerosis. Nat. Genet. 45, 1353–1360 (2013).

81. G. Trynka et al., Dense genotyping identifies and localizes multiple common and rare variant association signals in celiac disease. Nat. Genet. 43, 1193–1201 (2011).

82. J. Faraco et al., ImmunoChip study implicates antigen presentation to T cells in narcolepsy. PLoS Genet. 9, e1003270 (2013).

83. P. Morris et al., Large-scale association analysis provides insights into the genetic architecture and pathophysiology of type 2 diabetes. Nat. Genet. 44, 981–990 (2012).

84. V. Iotchkova et al., GARFIELD - GWAS Analysis of Regulatory or Functional Information Enrichment with LD correction. bioRxiv, 085738 (2016).

85. H. Wickham, ggplot2: Elegant Graphics for Data Analysis (Springer, 2009), Use R!

86. H. Li et al., The Sequence Alignment/Map format and SAMtools. Bioinformatics. 25, 2078–2079 (2009).

87. Heger, C. Webber, M. Goodson, C. P. Ponting, G. Lunter, GAT: a simulation framework for testing the association of genomic intervals. Bioinformatics. 29, 2046–2048 (2013).

